# Photomanipulation of minimal synthetic cells: area increase, softening and interleaflet coupling of membrane models doped with azobenzene-lipid photoswitches

**DOI:** 10.1101/2023.01.03.522478

**Authors:** Mina Aleksanyan, Andrea Grafmüller, Fucsia Crea, Vasil N. Georgiev, Naresh Yandrapalli, Stephan Block, Joachim Heberle, Rumiana Dimova

**Affiliations:** Max Planck Institute of Colloids and Interfaces, Science Park Golm, 14476 Potsdam, Germany; Institute for Chemistry and Biochemistry, Freie Universität Berlin, Berlin, Germany; Department of Physics, Freie Universität Berlin, Berlin, Germany

## Abstract

Light can effectively interrogate biological systems in a reversible and physiologically compatible manner with high spatiotemporal precision. Understanding the biophysics of photo-induced processes in bio-systems is crucial for achieving relevant clinical applications. Employing membranes doped with the photolipid azobenzene-phosphatidylcholine (azo-PC), we provide a holistic picture of light-triggered changes in membrane kinetics, morphology and material properties obtained from correlative studies on cell-sized vesicles, Langmuir monolayers, supported lipid bilayers and molecular dynamics simulations. Light-induced membrane area increase as high as ∼25% and a 10-fold decrease in the membrane bending rigidity is observed upon *trans*-to-*cis* azo-PC isomerization associated with membrane leaflet coupling and molecular curvature changes. Vesicle electrodeformation measurements and atomic force microscopy reveal that *trans* azo-PC bilayers are thicker than POPC bilayer but have higher specific membrane capacitance and dielectric constant suggesting an increased ability to store electric charges across the membrane. Lastly, incubating POPC vesicles with azo-PC solutions resulted in the insertion of azo-PC in the membrane enabling them to become photoresponsive. All these results demonstrate that light can be used to finely manipulate the shape, mechanical and electric properties of photolipid-doped minimal cell models and liposomal drug carriers, thus, presenting a promising therapeutic alternative for the repair of cellular disorders.

## 1. Introduction

Conversion of light into mechanical energy as established with photoresponsive molecules provides a clean and renewable source of energy for the fundamental units of life, which offers a potential solution to reducing the demand of rapidly-depleting natural resources as well as building a more sustainable world for future generations^[1]^. Examples include light-driven biocompatible micropumps/microrobots to generate fluid flow^[2]^ as well as transport of macromolecules^[3]^ in biotechnology applications, photoactive micro/nanomotors for wastewater treatment^[4]^ as well as heavy metal removal and sensing^[5]^, thus providing applications in environmental remediation^[6]^. One of the advantageous features of light-induced processes is the high spatiotemporal precision allowing control over the targeted system^[7]^. Adjustment of exposure time, intensity, and wavelength of irradiation also reduce the number and amount of byproducts since light-induced processes generally do not require additional reagents^[7b]^. Photoprocesses are usually fast and reversible, in which light is used to interconvert photochemically active materials, known as photoswitches, between the low energy, thermodynamically favorable state and high energy, kinetically favorable metastable states ^[8]^.

Among the myriad of photoresponsive molecules in the literature^[9]^, azobenzene-derived photoswitches are most commonly studied^[10]^ and the spectrum of applications includes molecular solar thermal energy storage^[1b, 11]^, catalysis of chemical reactions^[12]^, generation of photostructured polymers^[13]^, molecular recognition^[14]^, modulation of neurotransmission^[15]^, design of photochromic materials^[16]^, drug delivery systems^[17]^, optoelectronics^[18]^ and photopharmacological tools^[19]^. Typically, azobenzene derivatives isomerize from thermodynamically stable *trans* state to metastable *cis* isomer (π-π* transition) with the effect of UV-A illumination (365 nm) whereas the irradiation of blue light (465 nm) favors n-π* transition and reverses the process.^[20]^ Herein, we employ 1-stearoyl-2-[(E)-4-(4-((4-butylphenyl)diazenyl)phenyl)butanoyl]-sn-glycero-3-phosphocholine photoswitch to build a synthetic photobiomachinery to control membrane shape and mechanical properties of minimal cells for potential biomedical applications.

In biological cells, it is known that cellular morphology plays an important role in regulating cellular activities such as endocytosis^[21]^, exocytosis^[22]^, gene expression mechanisms^[23]^, stem cell differentiation^[24]^ and proliferation^[25]^. The abnormal changes in the membrane morphology (e.g. excess membrane area with respect to enclosed volume caused by external triggers) and the incompetency of biological cells to modulate their membrane properties can be associated with accompanying implications in cellular homeostasis^[26]^, pathological developments^[27]^, cancer progression^[28]^ or apoptosis^[29]^. A fast-responding external trigger such as light for controlling membrane area and mechanical properties can facilitate transmembrane transport and exchange of substances across the cell membrane thus reducing the above-mentioned harmful effects stemming from the malfunction of cellular processes.

The response of membranes to external triggers can be visualized in giant unilamellar vesicles (GUVs)^[30]^. GUVs are occasionally referred to as minimal cells, because of their size and features allowing membrane reconstitution and encapsulation of important cellular elements. Because of their large size, GUVs offer the possibility to directly monitor the membrane under a microscope. Light– triggered changes have been investigated on GUVs to interrogate, among others, (i) light-sensitive proteins embedded in the membrane such as the photoreceptor bacteriorhodopsin^[31]^, (ii) the photoactivation of channel proteins^[32]^, (iii) membrane-embedded fluorescent dyes which raise the membrane tension and can cause transient poration under irradiation^[33]^. The effect of azobenzene derivatives have also been studied with giant vesicles. Examples include light-triggered changes in membrane properties^[32b, 32c, 34]^ as well as phase state or fluidity^[34c, 34d, 35]^. However, some of the studies on membrane mechanical properties, area changes and thickness lack accurate and consistent characterization (as we discuss in section 4), while quantitative evaluation of the photoisomerization effects on other characteristics such as the membrane capacitance, dielectric constant and interleaflet coupling is still missing. Furthermore, to the best of our knowledge, the dose-dependent effect of azobenzene derivatives has been explored only to a very limited extent and mainly with water-soluble light switches, which upon insertion into the GUV membrane induce bursting or morphological transformations^[32c, 32d]^. Indeed, understanding the fraction-dependent effect of membrane photoswitches is important when considering potential implementation of these molecules for the local modulation of membrane characteristics such as thickness, tension, fluidity, and permeability. Water-soluble derivatives are probably less suitable for such applications than more hydrophobic membrane analogues.

In this work, we investigate the dose-dependent function of an azobenzene-derived lipid analogue (azobenzene-phosphatidylcholine, azo-PC, see Fig. 1A) applied to the lipid bilayer of giant vesicles to construct artificial photoswitchable cell mimetics from sustainable biomaterials. We subject this system to a thorough investigation for potential biomimetic purposes and applications in biomedical research. In parallel to our minimalistic cell model based on GUVs, we probe the response of Langmuir monolayers, large unilamellar vesicles (LUVs) and supported lipid bilayers (SLBs) with analogous composition and bilayer patches constructed with molecular dynamics (MD) simulations to interrogate the system at leaflet and molecular level, respectively. Characterizations of light-induced membrane shape transformations of GUVs membranes containing azo-PC have been reported previously^[34a, 34c, 34d]^, however, lacking the quantitative link between material properties and membrane parameters such as changes in area and thickness, morphology, elastic and electrical properties, and their relation to organization and restructuring at the molecular level. Herein, by combining a comprehensive set of experimental methods and model membrane systems with MD simulations, we provide a holistic picture of the photoresponse of membranes containing azo-PC. We first establish an experimentally undemanding approach for direct evaluation of photo-induced area changes based on GUV deformation in electric fields. The method is then employed to characterize the photoswitch isomerization kinetics and reversibility. Comparison across different model systems is also provided. We provide the first measurements on membrane thickness and specific capacitance of azo-PC bilayers and elucidate the interrelation between dynamics of photoswitching, membrane material properties and interleaflet coupling, changes in membrane area and thickness. Finally, we exogenously introduce azo-PC in preformed vesicles and quantitatively monitor the efficiency of photoswitching to test the potential applicability of this photoswitch in cellular studies. Understanding the underlying photoswitching dynamics on the membrane material properties can elucidate the light-controlled micromanipulation of cellular processes and optimization of light-triggered drug delivery platforms for potential applications of azo-PC containing bio-engineered minimal cells in photopharmacology.

**Figure 1:**
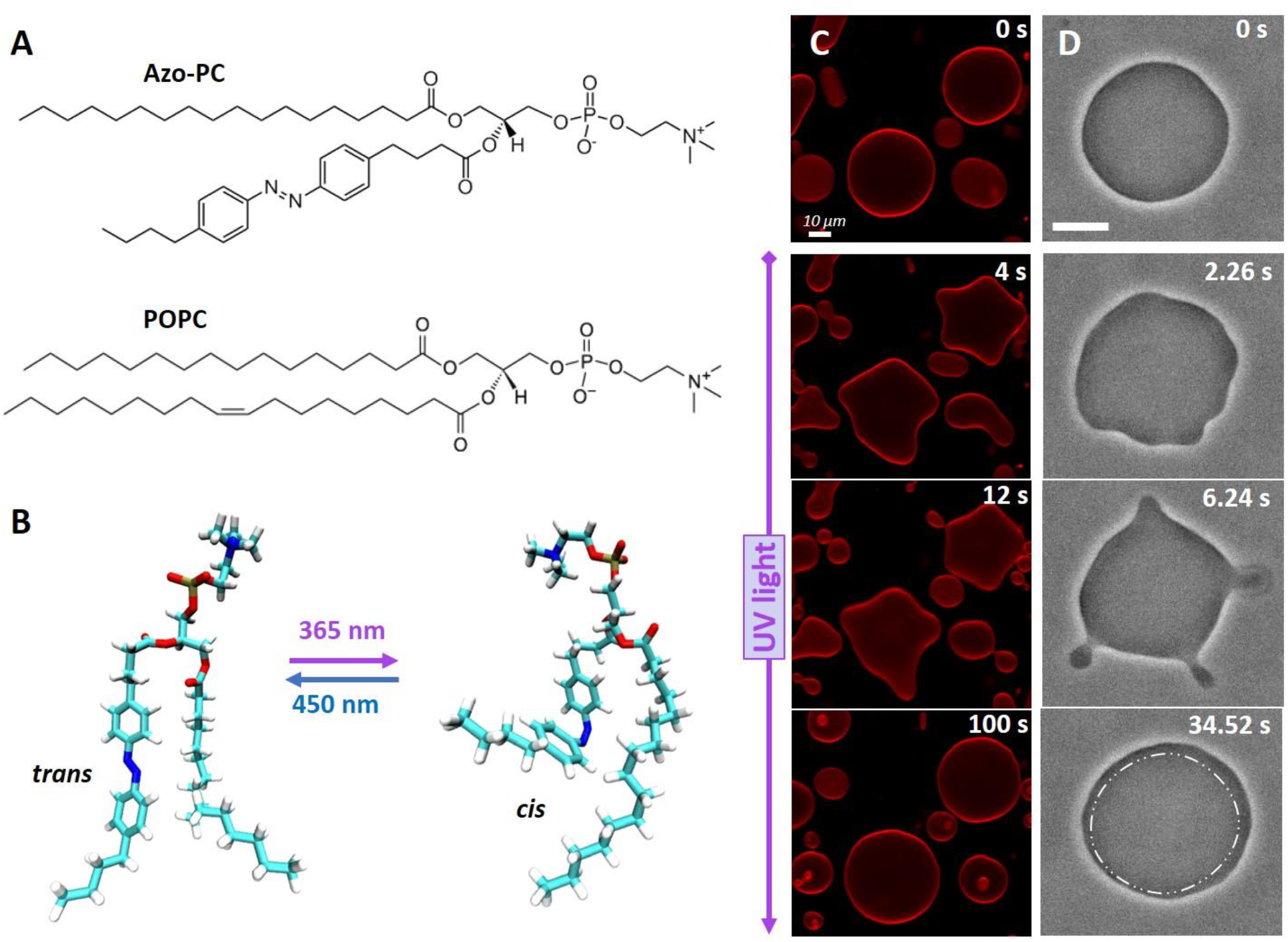
*Trans*-to-*cis* photoisomerization of azo-PC triggers vesicle shape changes and area increase. (A) Chemical structures of azo-PC and the POPC. (B) Representative snapshots of the molecular conformational changes upon photo isomerization of azo-PC obtained from MD simulations. (C) Confocal cross section images of 100 mol% azo-PC GUVs labelled with 0.1 mol% Atto-647N-DOPE monitored during photoisomerization. Upon UV irradiation (365 nm), the GUVs undergo complex shape transformations of outward budding and bud re-adsorption over time; the time stamps are shown in the upper part of the images. (D) Phase contrast microscopy showing a time sequence of the *trans*-to-*cis* photoisomerization response of 50 mol % azo-PC doped vesicles (azo-PC:POPC 50:50) under UV illumination, see also Movie S2. Budding and bud re-adsorption occur over time. The area of the vesicle increases: for comparison, the dash-dotted contour in the last image shows the approximate GUV contour before irradiation (first snapshot). Scale bars correspond to 10 μm.

## 2. Materials and methods

### 2.1. Vesicle preparation

GUVs were prepared by the electroformation method^[30a]^ at room temperature (23° C). Varying molar fractions of azo-PC (0, 5, 10, 25, 50, 100 mol %) and 1-palmitoyl-2-oleoyl-sn-glycero-3-phosphocholine (POPC) (both purchased as chloroform solutions from Avanti Polar Lipids, Alabaster, AL) were dissolved in chloroform to a concentration of 4 mM. Then, 8 μL of this lipid solution was spread as a thin film on a pair of indium-tin oxide (ITO)-coated glass plates (PGO GmbH, Iserlohn, Germany), which are electrically conductive. A stream of N_2_ was applied to evaporate most of the chloroform, and the plates were subsequently placed under vacuum for two hours to remove traces of the solvent. For chamber assembly, a Teflon spacer of 2 mm thickness was placed between the ITO-glass plates and the chamber was filled with a solution of 100 mM sucrose (Sigma Aldrich, St. Louis, USA) to hydrate the lipid film. For the GUV electrodeformation studies, the sucrose solution was also supplemented with 0.5 mM NaCl to ensure higher conductivity of the internal solution and thus prolate deformation of the GUVs^[36]^. Electroswelling was initiated by applying a sinusoidal AC electric field at 10 Hz frequency with a 1.6 V_pp_ (peak to peak) amplitude for 1 hour in the dark. GUVs were then transferred to light-protective glass vials for storage at room temperature and used the same day. For microscopy observations during electrodeformation studies, GUV solutions were diluted 8-fold with 105 mM glucose. The osmolarity was adjusted with an osmometer (Osmomat 3000, Gonotec GmbH, Germany). For bending rigidity measurements, GUV solutions were diluted 1:1 in 85 mM sucrose and 20 mM glucose to avoid gravity effects affecting the fluctuation spectra^[37]^. For confocal microscopy observations, GUVs were prepared from 100 mol% azo-PC further doped with 0.1 mol% 1,2-Dioleoyl-sn-glycero-3-phosphoethanolamine labeled with Atto 647N (Atto-647N-DOPE)(Avanti Polar Lipids).

For membrane capacitance measurements, following previously established protocols^[38]^, the GUVs were electroformed in 40 mM sucrose solution containing 0.3 mM NaCl at 50 Hz and 1.5 V_pp_ for 1 hour. Then, they were harvested and 9-fold diluted in 45 mM glucose solution containing 0.6 mM NaCl. The resulting conductivities of the GUV inner (42.70 μS/cm) and outer (81.67 μS/cm) solutions measured via Seven-Compact Conductivity Meter (Mettler Toledo, Ohio, USA) were thus adjusted to result in conductivity ratio of inner to outer solution Λ = 0.52.

Large unilamellar vesicles (LUVs) were prepared via extrusion at room temperature. Azo-PC (0, 50 and 100 mol %) and POPC (Avanti Polar Lipids, Alabaster, AL) were dissolved in chloroform to a concentration of 10 mg/mL. An aliquot of 100 μL of the lipid solution was dried under a gentle stream of nitrogen until the formation of a film on the walls of a glass vial. The vial was placed in a desiccator under vacuum overnight for the complete evaporation of the chloroform. The dried lipid film was hydrated with 1 mL Milli-Q water and agitated with a vortex mixer. The suspension was extruded 31 times through a polycarbonate membrane (Whatman® Nuclepore™ Track-Etched Membranes, Merck, Germany) with pore size of 200 nm using a mini-extruder (Avanti Polar Lipids).

### 2.2. Preparation of Langmuir monolayers and area change measurements in a Langmuir-Blodgett through

Azo-PC and POPC lipids were diluted to 1 mg/mL in chloroform. The two solutions were mixed in the desired ratios: 0 %, 10 %, 25 %, 50 %, and 100 % azo-PC molar fraction in POPC. An aliquot of lipid chloroform solution was deposited on the water surface of a commercial Langmuir-Blodgett trough (Kibron MicroTroughX, Kibron, Finland) with an available surface of 80 × 350 mm^2^, in the correct amount (about 16 μL) to yield an initial lipid density at the air-water interface of 100 Å^2^/lipid. A UV rod lamp (365 nm, Camag, Switzerland) and an array of blue LEDs (450 nm, Luxeonstar, Canada) placed above the film illuminated the trough surface with a power density of 30 μW cm^−2^ and 10 mW cm^-2^, respectively. The power density was measured at the sample position with a handheld power meter (LaserCheck, Coherent, USA). The film was first illuminated for 10 minutes with blue or UV light, respectively, to obtain either the *trans* or the *cis* isomer of the azo-PC component of the deposited lipids, and then compressed to a lateral pressure of 30 mN m^-1^ reducing the available area by compression with Teflon barriers at a speed of 35 mm min^-1^.

Compression isotherms during illuminations with both light sources and for all azo-PC molar fractions, were recorded with a sampling rate of 0.25 s^-1^. To calculate the area expansion as a percentage of the footprint of azo-PC in the *trans* state, the area per lipid at 30 mN/m occupied under blue-light illumination (azo-PC in the *trans* state) was subtracted by the value obtained under UV light illumination (azo-PC in the *cis* state) and divided by the first value. A subsequent new *trans* isotherm was recorded to then calculate the area reduction in analogy. For each azo-PC fraction, the results from four or five illumination cycles on different films were calculated.

### 2.3. Vesicle electrodeformation and area change measurements

Electrodeformation experiments to measure vesicle area changes^[39]^ were conducted in a commercial Eppendorf electrofusion chamber (Eppendorf, Germany) described previously. The chamber has two parallel cylindrical platinum electrodes (92 mm in radius) spaced 500 μm apart. GUVs between the electrodes were exposed to a sinusoidal AC field at 1 MHz with a 5 V (peak to peak) amplitude. In the absence of an electric field, the vesicles are quasi-spherical and exhibit visual fluctuations. The membrane area stored in these fluctuations is not possible to assess directly from the microscopy images. The mild AC-field deforms the GUVs and the type of the deformation depends on the field frequency and conductivity ratio between the internal and external GUV solutions^[40]^. At the conditions employed here, the GUVs adopt prolate ellipsoidal shapes, allowing for the precise measure of the total vesicle area from the vesicle geometry:

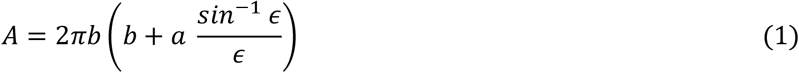

where *a* and *b* are the vesicle semi-axes along and perpendicular to the applied electric field, respectively, and *ϵ* is the ellipticity defined as *ϵ*^2^ = 1 − (*b*/*a*)^2^. Subsequently, the GUVs were irradiated with UV and blue light while the AC-field was still on and vesicles were recorded for 25-30 seconds at an acquisition speed of 8 frames per second (fps). Subtracting the initial vesicle area in the absence of irradiation (but with applied field), *A*_*i*_, from the vesicle area when exposed to light with specific wavelength, *A*_l_, yields the percentage of relative area increase as 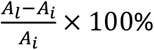 associated only with the photoisomerization of the azo-PC molecules.

The length of the semi-axes was measured from the recorded vesicle images either manually or using a home-developed software for GUV contour detection^[41]^. Data and statistics for GUVs with varying azo-PC fractions were plotted and analyzed using Origin Pro software. At least 10 GUVs from 3 separate sets of experiments for each investigated azo-PC fraction were used to plot the graphs. The statistical significance of the vesicle area changes due to photoswitching was tested with the one-way analysis of variance (ANOVA).

### 2.4. Vesicle imaging and irradiation

UV-induced shape transformations of 100 mol% azo-PC GUV was monitored through Leica TCS SP8 scanning confocal microscope (Wetzlar, Germany) using a HC PL FLUOTAR 40×/ Numerical Aperture (NA) 0.6 (air) objective. The pinhole size during the experiment was set to 1 AU (Airy units) and the scanning speed was 400 Hz in unidirectional mode. The Atto-647N-DOPE dye was excited with a HeNe 633 nm laser with 3 % (laser intensity) and the emission signal was collected with a HyD (hybrid) detector in the range 645-705 nm. In order to observe photoisomerization response of GUVs, the external 365 nm UV-LED was attached to the condenser of the confocal microscope. The observation chamber was made of two cover slips (22×40 mm^2^ and 22×22 mm^2^, Knittel Glass, Germany) sandwiching a spacer with a thickness of 1 mm.

Electrodeformation measurements were performed under phase contrast mode of an inverted microscope Axio Observer D1 (Zeiss, Germany), equipped with a Ph2 20× (NA 0.5) objective. Images were taken with an ORCA R2 CCD camera (Hamamatsu, Japan); see also section 2.5 for the setup used for bending rigidity measurements. The GUVs were placed in an Eppendorf electrofusion chamber with approximate thickness of 8 mm (other specifications are indicated in section 2.3). For UV and blue irradiation of the samples, the light from the microscope mercury lamp (HBO 100W) mounted in epi-illumination mode passed through 365 and 470/40 nm filters, respectively. The irradiation power of the HBO lamp was 60 mW cm^−2^ for the UV filter set (365 nm) and 26 mW cm^−2^ for the blue filter. Power intensities were measured with LaserCheck power meter after the objective and at the position of the sample.

### 2.5. Bending rigidity measurements

The membrane bending rigidity was measured with fluctuation spectroscopy of the thermal undulations of quasi-spherical vesicles as reported previously^[37, 41]^. Membrane fluctuations were observed under phase contrast of an inverted microscope Axio Observer D1 (Zeiss, Germany) equipped with a PH2 40 x (0.6 NA) objective. Sequences of 3000 images we recorded with Pco.Edge sCMOS camera (PCO AG, Kelheim, Germany) at an acquisition rate of 25 fps and exposure time of 200 μs (the same camera was used also for imaging kinetics of vesicle deformation under photoisomerization imaged at 100 fps). The vesicle contour was detected and analyzed with a home-developed software^[41]^. Low crossover modes were selected as 3 – 5 for eliminating the effects of vesicle tensions. Only defect-free, quasi-spherical vesicles with low tension values in the range 10^−7^– 10^−9^ N m^-1^ and 10 25 μm in radius were analyzed.

### 2.6. MD simulations

POPC lipids were modeled using the amber Lipid14 force field ^[42]^; parameters for the azo tail of azo-PC were taken from the optimized parameters for azobenzene from ref. ^[43]^ based on the general AMBER force field^[44]^. Figure 1B shows the simulated structures of azo-PC and the respective conformations under UV and blue light. The partial charges for the tails were derived following the methodology used in the Lipid14 force field ^[42]^ using 50 conformations from a 50 ns MD trajectory and the R.E.D. tool scripts ^[45]^. The topologies for azo-PC and POPC lipids were converted using the glycam2gmx.pl script^[46]^.

The initial topology of the POPC bilayer structure with 400 lipids was generated using the CHARMM-GUI ^[47]^ and charmmlipid2amber.py script ^[48]^; coordinates for the azo-PC bilayers were created from the POPC bilayer by fitting and replacing the required number of oleoyl tails with azo tails using VMD ^[49]^. The bilayer systems were then solvated with 24467 TIP3P water molecules ^[50]^.

All simulations were performed using GROMACS version 5.1.2^[51]^. Systems were energy minimized with steepest descend and equilibrated with and without position restraints on the lipids for a total of 21 ns using the weak coupling scheme ^[52]^ to relax the size of the simulation box. Production runs were performed for 100 ns at 303 K, applying the Nose-Hoover thermostat ^[53]^ and Parrinello-Rahman barostat ^[54]^ to keep the pressure and temperature constant.

The bilayer hydrophobic thickness was defined from the distance between the maxima in the phosphate atom distribution along the bilayer normal. Error estimates were calculated as the standard deviations from 8 subsets of the time frames from the trajectory. Bilayer elastic properties were calculated as follows: The bilayer stretching modulus *K*_*A*_ was obtained from simulations of all bilayer compositions at lateral pressures of -2, -4, -8, -12 and -16 bar. *K*_*A*_ is found from a linear fit of 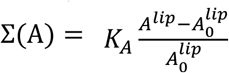, where ∑ is the mechanical tension, *A*^*lip*^ the area per lipid and 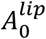 the area per lipid of a bilayer with zero mechanical tension. The standard deviation was calculated over all systems. The bending modulus κ was calculated from the lipid splay distribution as described in Ref. ^[55]^ using the python modules available at https://github.com/njohner/ost_pymodules/^[56]^.

The potential of mean force (PMF) for displacing phospholipid head-groups along the reaction coordinate *z* perpendicular to the lipid bilayer, was calculated by using umbrella sampling. For each lipid displaced across the membrane, 7 evenly spaced umbrella windows were created between *z* = 0 and *z* = 3 nm, with a harmonic umbrella potential with force constant 200 kJ mol^-1^ centered at that position. Two lipids, one from each monolayer, were restrained with an offset of 3 nm between their potentials. In this way, at *z* = 0 for one lipid, the second lipid had *z* = 3 nm. Each umbrella window was simulated for a total of 200 ns, where the first 50 ns were discarded as equilibration time. The PMF profiles were then constructed using the weighted histogram analysis method^[57]^. Error bars reflect the differences between the PMFs of the separate lipids.

### 2.7. LUV area change measurements via dynamic light scattering (DLS)

The average diameter of the LUVs was measured through Zetasizer Nano ZS90 DLS (Malvern Instruments, Malvern, United Kingdom) equipped with a 632.8 nm 4mW HeNe laser and measuring the scattered light at 173°. In order to detect the UV-induced area change of the LUVs, the UV LED (used also for the measurements with GUVs) was mounted inside the measuring compartment of the Zetasizer. Three replicates were produced for each light condition for each sample. The average size distribution of the LUVs was plotted and the area of LUVs was calculated from the formula of a sphere, 4πR^2^, in which R is the LUV radius. Area increase is defined similarly to that for the GUV measurements, namely as 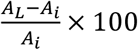 where *A*_*L*_ here is the LUV area after the UV exposure and *A*_*i*_ is the initial area of the vesicle before the UV irradiation.

### 2.8. Preparation of supported lipid bilayers (SLBs) and measurement of bilayer thickness through atomic force microscopy

Supported lipid bilayers (SLBs) were prepared based on a previously reported protocol^[58]^ with some modifications. Briefly, a chloroform solution of POPC or azo-PC lipids was dried in a clean glass vial under nitrogen flow and in a desiccator for 45 min. A 1 mg/mL final concentration (approx. 1 mM) of multilamellar vesicle suspension was produced by adding buffer (10 mM sodium citrate, 100 mM NaCl and 0.5 mM Ethyleneglycol-bis(β-aminoethyl)-N,N,N’,N’-tetraacetic Acid (EGTA), pH 4.5, 45 °C; all materials from Merck, Germany) to the dried lipid layer after overnight incubation at room temperature. The lipid suspension was subjected to bath sonication for 30 min before performing 21 extrusions through a 50 nm membrane using the mini extruder at 45 °C. Thus obtained small unilamellar vesicles (SUVs) were used to produce supported lipid bilayers on a freshly cleaned (first with 5% sodium dodecyl sulfate solution, then subjected for 1h and 20 min with freshly prepared piranha solution (H_2_SO_4_:H_2_O_2_ ratio of 3:1) under bath sonication) petri dish with glass bottom (μ-Dish, ibidi GmbH). After thoroughly rinsing with deionized water, 400 μL of SUV suspension (2 μM) was pipetted into the petri dish and incubated for 20 min at 45 °C. After a rinsing step (to remove excess SUVs), the SLBs were used for AFM measurements at room temperature.

SLB thickness measurements were performed using a JPK NanoWizard 3 (Bruker Nano GmbH, Berlin, Germany) atomic force microscope (AFM) mounted on an inverted microscope (model IX3-CBH, Olympus Corp., Japan). The AFM was equipped with a SNL-10 cantilever (Bruker AFM probes; spring constant: 0.35 N m^-1^, tip height: 7 μm, nominal tip radius: 2 nm; resonance frequency: ∼89 kHz in air). AFM measurements were performed using QI™ mode, yielding images of 256 pixels × 256 pixels. The images were taken at a vertical speed of 89 μm/s, z-length of 300 nm, and a cycle period of 3.4 ms per pixel. The SLBs produced in the petri dish were directly used for measurement without further sample treatment. In the case of SLBs containing azo-PC lipids, *trans* to *cis* transition was achieved using the DAPI filter (350/50 nm wavelength) from source SOLA light engine (Lumencore®, USA) of the inverted microscope (with a typical output of ∼5 mW cm^-2^) under dark conditions.

The preparation conditions described above typically resulted in samples, in which patches of SLBs were distributed across the glass substrate. The corresponding AFM images were analyzed using home-written scripts implemented in MatLab (MathWorks, Natick, MA)^[59]^. The open source software Gwyddion^[60]^ was used to store the height channel of the raw AFM data files as column-separated value (CSV) files, which can be directly loaded into MatLab. In each height map, the substrate (i.e., the glass surface) was assigned to a height value of 0 by fitting and subtracting a polynomial of first order to and from each line of the height map. Pixels, which are not related to the substrate (e.g., SLB-related pixels), were automatically rejected from this process using the following iterative procedure: (i) line-wise linear fit to all pixels, (ii) rejecting all pixels being higher than 1 nm from the result of the first fit, followed by a line-wise linear fit to all remaining pixels, (iii) rejecting all pixels being higher than 0.5 nm from the result of the second fit, followed by a line-wise linear fit to all remaining pixels. Afterwards, the height maps were smoothed using a 10 × 10 pixel^2^ moving average. The thickness of SLB patches was determined by cropping patch-containing areas from the AFM height map and by generating histograms of the height values of all pixels from such crops. These histograms typically exhibited two prominent peaks, which correspond to pixels of the substrate (centered around a height value of 0) and of the SLB patch, respectively; see e.g. Fig. S7B, left column). The height distance of these peaks measures the thickness of the corresponding SLB patch.

### 2.9. Specific membrane capacitance measurements and evaluation of the membrane dielectric constant

Membrane capacitance of pure POPC and 100 mol% *trans* azo-PC GUVs were deduced using an established protocol based on vesicle electrodeformation^[38a, 61]^. The field frequency is gradually varied between 500 Hz and 1 MHz at 10 kV/m field strength. In this frequency range, the GUVs can adopt different morphologies (prolate, oblate or spherical) depending on the conductivity ratio 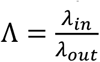 between the inner and outer GUV solutions^[36, 40a]^. For Λ < 1, the GUVs adopt prolate shapes at low field frequencies. With increasing frequency, the aspect ratio *a*/*b* decreases and at a critical frequency *f*_*c*_, the vesicle shape is a sphere 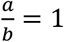. Further increase in the frequency leads to oblate morphologies, 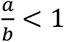. Critical frequency depends on the specific membrane capacitance, *C*_*m*_ ^[62]^:

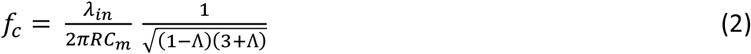

where *R* is the GUV radius. The critical frequency can be obtained from frequency-sweep experiments in AC field. The GUV solution (see 2.1 for preparation) was transferred into commercially available Eppendorf electrofusion chamber described in section 2.3 and exposed to AC field. GUVs were monitored via Axio Observer D1 Phase contrast microscope equipped with Ph2 20× (NA 0.5) objective. Images were acquired with ORCA R2 CCD camera (Hamamatsu, Japan) and the aspect ratio changes of GUVs were analyzed from the contour of the vesicles via lab-owned software^[41]^. For the statistics, 10 GUVs were analyzed with radii varying between 3 μm and 10 μm for each composition. The specific membrane capacitance was deduced from the slope of a linear fit to the critical frequency as a function of 1/*R* using Eq. 2.

The experimentally measured capacitance *C*_*m*_ is the resultant of the capacitances of bare lipid bilayer, *C*_*B*_ and of the ionic double layers in the inner and outer solutions *C*_*D,in*_ and *C*_*D,out*_ so that

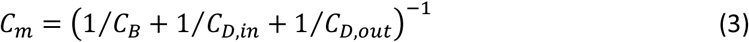

The membrane, can be considered as a two-dimensional surface with dielectric permittivity *ε*_*r,B*_*ε*_0_, where *ε*_*r,B*_ is the relative (dimensionless) dielectric permittivity of the bilayer and *ε*_0_ is the vacuum permittivity, *ε*_0_ ≈ 8.85 × 10^−12^ F/m. The dielectric permittivity of the bilayer relates to its capacitance and thickness *d* as:

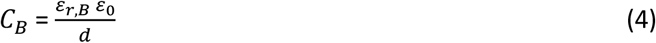

The capacitive contributions *C*_*D,in*_ and *C*_*D,out*_ can be estimated from the dielectric constant of the water solution *ε*_*r,W*_ ≈ 80 and the thickness of the Debye length, *λ*_*D*_. For the inner and outer vesicle solutions used here *λ*_*D*_ is 17.5 nm and 12.4 nm, respectively, yielding *C*_*D,in*_ = 4.05 μF/cm^2^ and *C*_*D,out*_ = 5.72 μF/cm^2^. These values and *C*_*m*_ were used in Eq. 3 to estimate the specific bilayer capacitance *C*_*B*_ for pure POPC and 100 mol% *trans* azo-PC membrane. Then, the bilayer dielectric constant *ε*_*r,B*_ was deduced from Eq. 4 by using membrane thickness values from the AFM studies in section 2.8.

### 2.10. Exogenous addition of azo-PC

For exposing preformed GUVs to exogenous azo-PC, pure POPC GUVs prepared in 100 mM sucrose solution were diluted 1:1 in 105 mM glucose solution to a final volume of 174 μL, which was then mixed with 1 μL of 2.72 mM azo-PC dissolved in 2:1 (vol) dichloromethane/methanol solution. The final concentration of azo-PC in the GUV suspension was 15.56 μM being close to the total amount of POPC lipids forming the vesicles, i.e. the final azo-PC-to-POPC ratio was about 1:1. After 20 minutes of incubation allowing the evaporation of the organic solvents, POPC GUVs enriched with azo-PC were placed in an observation chamber and exposed to UV and blue irradiation, respectively, and monitored by phase-contrast imaging.

## 3. Results

### 3.1. Vesicle shape deformation under light

Electroformation of pure azo-PC vesicles has been previously reported^[34a, 34c, 34d, 63]^, using relatively long swelling time (2-3 hours), and harsh conditions of high temperature (70 °C) and high voltage (10 Hz, 3–5 V) conditions (note that at such conditions double bonds of lipids may oxidize). Here, we reduced the GUV preparation time (to 1 hour) using much milder conditions, namely field strength of up to 1 V and room temperature. Pure azo-PC GUVs have been explored under dark-field and epifluorescence microscopy, to detect effects of illumination ^[34a, 34c]^. These approaches require intense white light illumination and a substantial fraction of a membrane fluorophore. Here, we implemented confocal microscopy, which offers higher resolution of the photo-induced membrane deformations and, compared to epifluorescence observations, requires only one-tenth of the fraction of the membrane fluorophore (0.1 mol%) for visualization. It is noted that high fractions of fluorophores affect membrane material properties such as bending rigidity^[64]^ and can cause oxidation and changes in membrane composition^[65]^. In addition, we probed the response of vesicles devoid of fluorescent dye using phase-contrast microscopy to eliminate potential dye effects. The UV irradiation in our confocal setup was implemented with an external source mounted at the microscope condenser (see Methods section 2.4). GUVs containing azo-PC and labeled with 0.1 mol % Atto-647N-DOPE were exposed to UV light to initiate *trans*-to-*cis* photoisomerization and observed for a few seconds. Before UV irradiation, GUVs were mostly defect free (at least 90 % of the population) exhibiting thermal fluctuations visible both in confocal and phase-contrast microscopy. Upon UV illumination, GUVs containing substantial fractions of azo-PC (50 or 100 mol%) undergo large shape transformations including complex budding events within a few seconds visible in confocal and phase contrast microscopy (Figs. 1C,D, S1 and Movies S1, S2 and S3). The vesicles increase in size (see last snapshot in Fig. 1D), but a quantitative assessment of the membrane area change is not feasible because of the unknown GUV geometry. Even if the vesicle appears as a sphere in the projected image, it can be flattened due to gravity into an oblate shape which has similar appearance in the images.

### 3.2. Assessing the light-induced membrane area change by GUVs electrodeformation, Langmuir monolayer isotherms, LUVs and MD simulations

To quantitatively characterize the membrane area change associated with photoisomerization, we employed GUV electrodeformation^[39a, 40b]^. In this approach, an alternating current (AC) field is applied before exposing GUVs to the UV light. Moderate strengths of electric fields are able to pull the excess area stored in thermal fluctuations. The vesicles deform into prolate or oblate shapes depending on the AC field frequency and conductivity ratio between the internal and external GUV solutions^[36, 40b]^. Due to gravity, oblate vesicles lie flat in the observation chamber and appear as spherical in the projected images not allowing access to their short semi-axis. On the contrary, prolate deformations, whereby the vesicle elongates along the field direction parallel to the bottom of the observation chamber, allow measuring both semi-axes *a* and *b* (Fig. 2A), and thus, the correct evaluation of the vesicle area. To induce prolate deformation, we prepared the GUVs in solutions containing salt (0.5 mM NaCl and 100 mM sucrose) and diluted the harvested vesicles in salt-free glucose solution (105mM). These conditions ensure higher conductivity in the GUV interior rendering them prolate under the applied AC field. Additionally, due to the refractive index differences between the interior and exterior sugar solutions, the GUVs appeared with a sharp contour under phase contrast observations (Fig. 1D) facilitating image analysis and area measurements.

**Figure 2:**
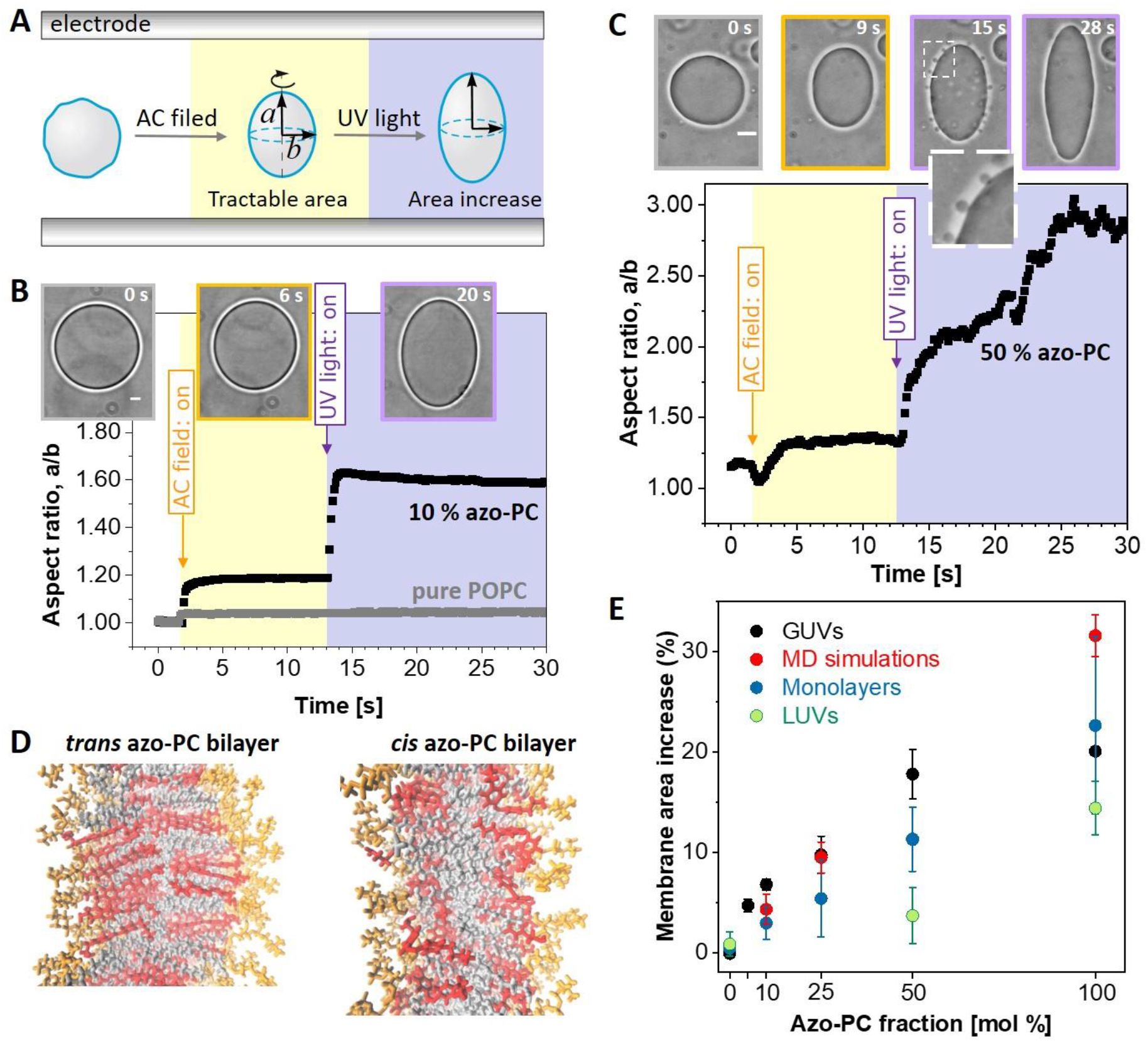
Area increase of membranes and monolayers doped with azo-PC when exposed to UV irradiation. (A) Sketch of the approach of GUV electrodeformation to assess the vesicle area change induced by UV light. The vesicles are first exposed to AC field (5 kV m^-1^ and 1 MHz) to pull out thermal fluctuations and deform them into a prolate ellipsoid with semi axes a and b. Then, while keeping the AC field on, the UV irradiation (365 nm) is initiated. (B, C) Electrodeformation and irradiation of GUVs made of pure POPC (gray trace in panel B) and containing 10 and 50 mol% azo-PC, see also Movies S4-S6 showing the response of these three vesicles. The snapshots show example images of the vesicles before applying the AC field (gray frame), after the application of AC field (orange frame) and when exposed to UV light (purple frame). A zoomed-up vesicle segment (dashed region) is given in C, showing the produced vesicle buds right after irradiation. The vesicle semi axes are used to calculate the vesicle area. Scale bars are 10 μm. (D) Snapshots from MD simulations bilayers composed of 100 mol% azo-PC in *trans* and *cis* conformation. The head groups of the lipids are in orange, the azo-benzene moiety in red, and the oleoyl tails in gray. The area of the bilayer increases and tits thickness decreases. (E) Membrane area expansion as assessed from GUV electrodeformation (black data show mean and standard deviations, SD; see Fig. S3 for data from individual GUV measurements), MD simulations (red), Langmuir monolayer isotherms (dark blue; see Fig. S3 for data from individual measurements) and LUVs measured with DLS (green). The LUV data is based on vesicle hydrodynamic radius leading to systematic underestimate for the area increase as UV-triggered morphological transitions (as those shown in panel C and Fig. 1C,D) cannot be accounted for.

We explored the area change in POPC vesicles containing 0, 5, 10, 25, 50 and 100 mol% azo-PC. The vesicles were first exposed to an AC field and the area was measured. Then, while keeping the AC field on, they were irradiated with UV light and the changes in vesicle shape were characterized in terms of changes in the vesicle aspect ratio *a*/*b*. The morphology change of each GUV was monitored over time under phase-contrast microscopy during the application of electric field and UV-light (see Fig. 2A-C, Fig. S2 and Movies S4-S6). The vesicle response to AC field is fast and completes within less than a second (the dynamics practically depends on the field strength and membrane viscosity^[66]^). The degree of deformation of the vesicles in the AC field in the absence of UV light showed variations from vesicle to vesicle. These are imposed by the vesicle size (affecting the magnitude of the Maxwell stress tensor deforming the vesicle) and the initial available excess area for deformation (which cannot be controlled as the GUV preparation method yields vesicles with different tensions).

UV illumination was typically applied ∼10 seconds after applying the AC field. Pure POPC GUVs did not show any response to UV light (see gray trace in Fig. 2B and Movie S4). The response of azo-PC-doped GUVs to UV light was very fast. GUVs containing 5, 10 and 25 mol% azo-PC reached their maximum deformation within a second after switching the UV light on (Figs. 2B, S2A and Movie S5). Increasing fractions of azo-PC resulted in larger membrane deformations in the form of buds (a couple of micrometers in size, see zoomed image in Fig. 2C) requiring longer times for the created membrane area to be pulled out into ellipsoidal shape. For GUVs with 50 mol% azo-PC, the buds pulled back by the electric field within roughly 12 – 15 seconds after applying the UV light, contributing to the vesicle elongation. These results indicate that budding and bud re-adsorption slowed down the deformation processes of GUVs containing high fractions of azo-PC (50 mol% and more), (see Figs. 2C, S2B and Movie S6). Indeed, vesicles made of 100 mol% azo-PC often did not reach perfect elliptical shapes affecting our accuracy for assessing the membrane area increase.

The area increase of the cell mimetic vesicles resulting from azo-PC photoswitching was calculated from the ellipsoid surface area of the GUVs at their maximal deformation (Eq. 1) and by subtracting the initial electric field-driven deformation in the absence of UV light. This subtraction eliminates effects associated with the initial membrane tension and the applied electric field. To account for the different vesicle sizes, we normalized the area by the initial one under electrodeformation in the dark. At least 10 vesicles per composition were examined. In the absence of azo-PC (pure POPC membranes), no detectable change in the vesicle area due to UV light was observed suggesting that the illumination conditions (intensity and duration) do not alter the membrane. However, with raising the molar fraction of azo-PC lipids in the membrane, GUVs area increase could rise up to 20 % (gray and black data in Fig. 2E). Similar but significantly smaller area change was found from dynamic light scattering (DLS) measurements on LUVs (see green data in Figs. 2E and S3, Methods section 2.7). It is worthwhile noting that the LUV hydrodynamic radius measured with DLS can be used to obtain only an apparent area change because an assumption for the vesicle shape (typically a sphere) is required. Thus, LUV measurements (as previously used in ref. ^[34a]^) do not properly represent the vesicle area increase as they do not account for morphological changes as those shown e.g. in Fig. 1C,D. Instead, DLS data leads to systematic underestimate (Figs. 2E). Thus, our results emphasize the superiority of GUVs as minimal cell model over LUVs for measuring area changes.

For higher fractions of azo-PC, the data on GUV area increase under UV exhibit larger scatter (larger standard deviations) and above 50 mol% azo-PC appear to reach saturation (Figs. 2E and S3). This is mostly due to the slow re-adsorption of the light-triggered buds as well as to the strongly elongated GUV shapes (observed for 100 mol% azo-PC) to which the elliptical approximation does not fully apply. Furthermore, the area of the photolipids is expected to be more packed and closely aligned at high azo-PC fractions, which could result in stronger dipole-dipole interactions between the azobenzenes in the lipid tails, potentially leading to photolipid clustering.^[34c, 67]^

In order to test these hypotheses, find out whether they are universal and not constraint to our minimalistic model system, and to gain deeper insight at molecular level into this photoswitchable minimal cellular system, we performed MD simulations of membranes with azo-PC in *cis* and *trans* state (Fig. 2D). We also examined Langmuir monolayers with different compositions exposed to UV and blue light, respectively (Fig. S4). Both MD simulations and monolayer isotherms yield excellent agreement with the data from our minimal cell model showing relatively linear increase of the light-induced area change with increasing azo-PC fractions in the membrane (Fig. 2E). Even though linear, the expansion data of the monolayers lie somewhat lower compared to that from the bilayer systems (MD and GUV membranes). This is to be expected as the monolayer lacks all inter-leaflet interactions that are present in the bilayer (and discussed in more detail below) and faces air as less similar environment compared to that of the lipid chain moieties. The MD simulation, consistent with monolayer data, reveal that the area per lipid decreases with increasing fraction of *trans* azo-PC in the membrane and the opposite is true for the *cis* azo-PC conformation. The combined effect of these opposite trends yields an increase in the bilayer area by up to ∼30 % for the pure azo-PC membrane (Fig. 2E). The quantitative match between the GUV model and MD simulations (except for the 100 mol% azo-PC case where the GUV electrodeformation approach lacks high accuracy) also suggests that under the selected irradiation conditions, full photoconversion of the azo-PC molecules occurs.

The simulations offer further insight in the origin of the membrane area changes as a function of photolipid fraction and isomerization state. The fairly rigid planar *trans* tails tend to orient along the membrane normal and can stack flatly against each other. The bent *cis* tails on the other hand orient more along the membrane plane and localize predominantly close to the headgroup-tail interface. As a result, the palmitoyl tails fill the region near the bilayer center, as clearly visible in the density profiles (Fig. S5).

### 3.3. Reversibility and kinetics of membrane response

To further examine the photoswitching efficiency on the bilayer, we investigated the reversibility and relaxation kinetics of the membrane response of azo-PC containing GUVs. Establishing a fully reversible and reproducible photoswitching process is an important criterion for the efficient regulation of membrane shape and material properties. In order to fully reverse the photoswitching from *cis*-to-*trans* isomerization, we applied blue irradiation after exposing azo-PC GUVs to UV light. Upon reversible and complete photoswitching, the area expansion of the vesicle due to *trans*-to-*cis* isomerization under UV light should be fully recovered and equal to the area shrinkage resulting from the *cis*–to-*trans* isomerization under the blue light. We compared the area changes under these two illumination conditions for GUVs containing 10 and 25 mol % azo-PC, where no complex budding events are observed and the area changes can be measured at high precision. The data are presented in Fig. 3A. Based on the results of the statistical tests, no significant differences were observed between the means of vesicle area changes for *trans*-to-*cis* and *cis*-to-*trans* isomerization for fixed membrane composition, indicating that light-induced morphological changes of azo-PC GUVs are reversible.

**Figure 3:**
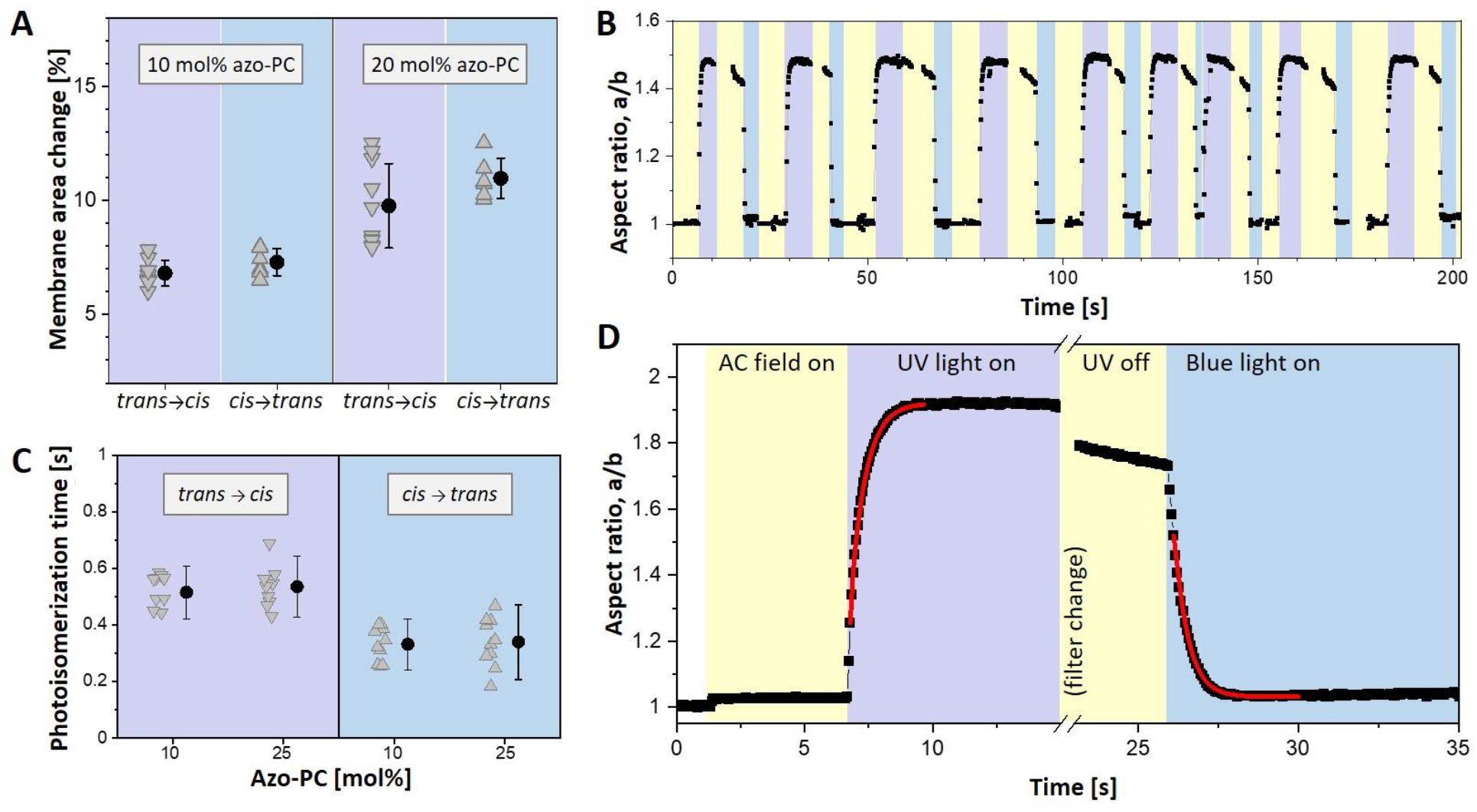
Photoswitching reversibility and kinetics assessed from the response of azo-PC GUVs exposed to UV and blue light. (A) Membrane area change measured on vesicles containing 10 and 25 mol % of azo-PC. *Trans*-to-*cis* isomerization upon UV illumination leads to area changes similar to that observed upon *cis*-to-*trans* isomerization under blue light. Each triangle indicates a measurement of an individual GUV. Mean and standard deviation values are also shown on the right. ANOVA test for null hypothesis testing for 10 and 25 mol % azo-PC GUVs gives respectively p = 0.136 and p = 0.065, indicating statistically insignificant difference for the *trans*-to-*cis* vs. *cis*-to-*trans* area change of membranes of a fixed fraction of azo-PC. (B) Multiple photoswitching cycles of 10 mol% azo-PC vesicle shown in terms of the degree of deformation (a/b, aspect ratio) under UV light (purple regions) and blue light (blue regions) sequentially switched on and off; the same vesicle is shown in Movie S7. Throughout the experiment, the GUV is continuously exposed to AC-field (5 kV.m^-1^ and 1 MHz; yellow). Purple and blue regions in the graph schematically illustrate the time intervals when UV and blue light are switched on. (C) Photoisomerization kinetics of 10 and 25 mol % azo-PC containing GUVs. Data from individual GUVs are shown with triangles (10 vesicles per composition and condition were measured). Solid circles and line bars show means and standard deviations. (D) Kinetic trace of the aspect ratio response to UV and blue light irradiation of a GUV containing 25 mol% azo-PC. The exponential fits (red curves) yield the respective time constants as plotted in panel C. A short period of time is needed to mechanically change the filter at the microscope turret, during which the recording of the vesicle is paused.

Similarly, the reversibility of swelling and shrinkage of GUVs due to photoswitching under UV and blue light were monitored several times (Fig. 3B, Movie S7). Vesicle deformation was fully reversible and could be switched back and forth over multiple cycles. Furthermore, we observed that the sharp contrast resulting from sugar asymmetry between GUV interior and exterior solutions was preserved, suggesting that during the multiple photoswitching cycles the membrane remains intact, i.e. the photoisomerization process did not generate any permeation or leakage over time. All these data illustrate that photoswitching under the selected irradiation conditions can be repeated without any sign of decomposition (of either azo-PC and POPC) or membrane leakage.

Based on the area swelling and shrinkage of electrodeformed GUVs under UV and blue light, we assessed the rates of isomerization from the response of GUVs containing 10 and 25 mol % azo-PC over the course of photoswitching. An example kinetic trace and the rates of photoswitching obtained from exponential fits to the data for *trans*-to-*cis* and *cis*-to-*trans* isomerization are shown in Fig. 3C, D. Our results demonstrated that the *cis*-to-*trans* exponential time constant (with a mean value of 335 ms) is shorter than the *trans*-to-*cis* response time (525 ms). This faster *cis*-to-*trans* photoswitching kinetics is understandable considering that the *trans* isomer is thermodynamically more stable. We note that these rates depend not on the molecular isomerization kinetics which are in the femtosecond to picosecond time range^[68]^ but is determined by the vesicle hydrodynamics and membrane viscosity. Photoisomerization kinetics did not show any difference between 10 to 25 mol % azo-PC containing vesicles.

### 3.4. Membrane mechanical properties

Considering the differences detected in the bilayer area, structure, and photoisomerization kinetics of *cis* and *trans* azo-PC GUVs, we hypothesized that membrane mechanical properties should also show differences depending on isomerization state and photoswitch fraction. We employed fluctuation spectroscopy^[37, 41, 69]^ as a contactless approach to characterize the bending rigidity of POPC membranes with various fractions of azo-PC in different isomerization states (Fig. S6A). In parallel, computational studies have been performed to calculate the bending rigidity of the simulated membranes from the real space fluctuations of the tilt and splay of lipid tails^[55]^. The absolute values obtained from experiment and simulations are expected to differ (as we find, see Fig. S6B), because the bending rigidity is sensitive to the composition of solutes in the bathing medium^[70]^ (pure water in simulations and sugar solutions in the experiment). To allow comparison, the results were normalized by the mean value of the bending rigidity measured for pure POPC membranes, Fig. 4A.

**Figure 4:**
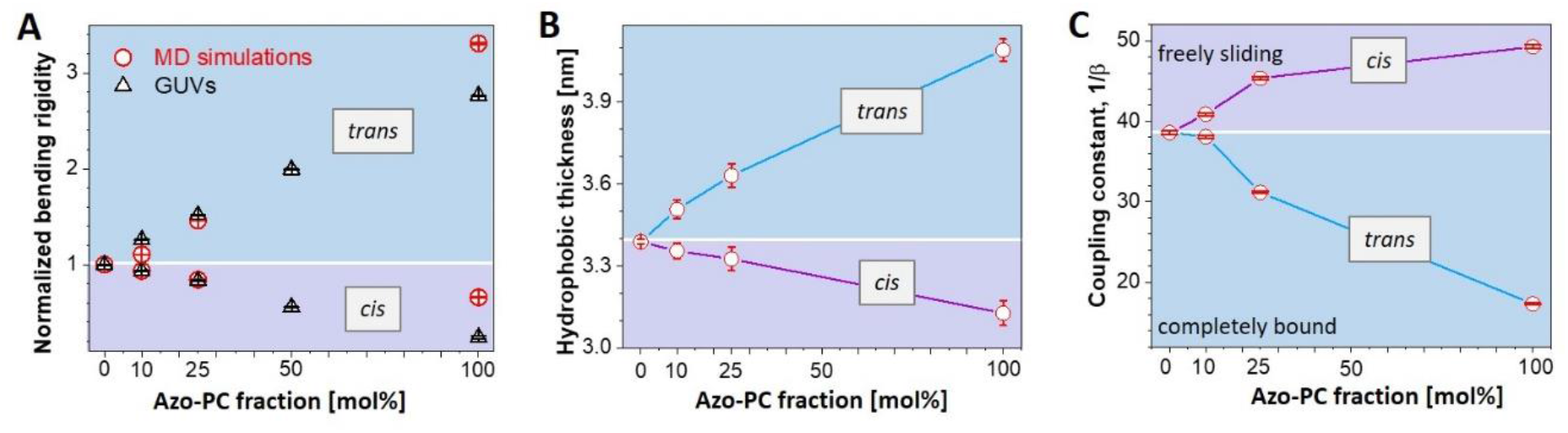
Bending rigidity, thickness and interleaflet coupling in membranes with various fractions of azo-PC in the *cis* and *trans* states. (A) Bending rigidity obtained from fluctuation spectroscopy (open triangles) and MD simulations (open circles). The results are normalized by the bending rigidity value of pure POPC (see non-normalized data in Figure S6). Blue and purple data correspond to *trans* and *cis* azo-PC, respectively. For each composition 10 GUVs are analyzed. Standard deviations are illustrated with line bars, smaller than the sizes of the symbols. (B) Bilayer thickness data at various fraction of azo-PC in *trans* and *cis* conformation obtained from MD simulations. (C) Interleaflet coupling in *cis* and *trans* azo-PC containing bilayers. The coupling constant is deduced from simulation data by using the formula based on polymer brush model^[71]^, in which the elasticity ratio scales quadratically with hydrophobic thickness of the bilayer (κ/*K* = *βd*^2^) and 1/β describes the coupling between the bilayer leaflets. Purple and blue trends demonstrate the coupling constants for *cis* and *trans* bilayers, respectively. Line bars are standard deviations and smaller than the size of the symbols.

We find excellent agreement between experiment and simulations demonstrating adeptness of the used force fields. Membranes doped with azo-PC in the *cis* conformation have lower bending rigidity compared to pure POPC and *trans* azo-PC GUVs, therefore, *cis*-photoisomerization of azo-PC softens the membrane. On the one hand, as the *cis* azo-PC fraction in the membrane increased from 0 to 100 mol%, the bending rigidity decreased 4 fold dropping down to values as low as 5 k_B_T (Fig. S6A); similar bending rigidity decrease has been observed upon the insertion of fusion peptides^[72]^ pointing to the destabilizing potential of azo-PC. On the other hand, equivalently increasing *trans* azo-PC fractions in the membrane stiffens the membrane 3 fold reaching bending rigidity values around 70 k_B_T (Figs. 4A and S6A); such bending rigidities are characteristic of membranes in the liquid ordered phase^[41, 73]^. This is indeed consistent with the strong alignment of the azo-PC tails structurally resembling liquid ordered phases.

A simplistic reason for the changes in the bending rigidity could be sought in changes in the membrane thickness due to photoisomerization. X-ray scattering studies on pure azo-PC vesicles have shown thinning of the bilayer by approximately 4-5 Å resulting from *trans*-to-*cis* isomerization^[63]^. Thinner membranes are generally softer and *vice versa* but such a small thickness change cannot account for the large bending rigidity changes we observe. We thus questioned the reported data and measured the thickness of POPC and azo-PC membranes from AFM on supported lipid bilayer patches (see Fig. S7 and Methods section 2.8). The thickness of POPC bilayers was measured as 4.7 ± 0.3 nm, which is in very good agreement with literature data^[74]^. In the case of 100 mol% azo-PC containing bilayer, thickness values of 6.2 ± 0.4 nm and 4.7 ± 0.4 nm were measured for the *trans* and *cis* conformations, respectively. This thickness change is more consistent with the membrane softening as shown from the bending rigidity measurements.

Considering that the MD simulations correctly represent the experimental findings on area and bending rigidity changes, we also explored how the membrane hydrophobic thickness varies with azo-PC fraction and isomerization. MD simulations show that isomerization and increasing azo-PC fractions can alter membrane thickness by almost 1 nm (Fig. 4B), consistent with the whole-bilayer thickness changes measured with AFM. The thickness of membranes containing azo-PC in the *trans* state are more influenced by the photolipid fraction.

In addition to thickness changes as a reason for softening the membrane, the bending rigidity behavior in Fig. 4A could also be related to dipole-dipole coupling between azobenzene groups in the acyl chains of the photolipids.^[34c, 67]^ Stronger molecular interactions between photolipids might lead to more densely packed bilayer giving less flexibility to the membrane for bending. Another factor modulating the membrane mechanical properties is the interleaflet coupling, which relates the bending rigidity, κ, the stretching elasticity, *K*, and the membrane thickness, *d*. The relation of the two elastic moduli κ and *K* has been theoretically considered^[71, 75]^ and experimentally explored for pure lipid membranes^[75a]^. Their ratio scales quadratically with the membrane thickness: κ/*K* = *βd*^2^, where the proportionality constant *β* described the coupling of the monolayers constituting the membrane. For 1/*β* = 12, the leaflets are completely bound^[75a]^, for 1/*β* = 48, they are unbound and freely sliding^[75b]^, while the polymer brush model^[71]^ predicts the intermediate value of 1/*β* = 24 for lipid bilayers. To resolve the leaflet coupling in our azo-PC membranes, we first assessed the stretching elasticity modulus *K* from MD simulations. This was achieved from fitting the tension versus area curves (see Methods section 2.6), which did not show any systematic dependence on azo-PC fraction in the bilayer. A combined fit to all data points resulted in a value of *K* = 221.8 ± 6.8 mN/m which we kept constant for all calculations. The coupling constant values were then estimated for the corresponding fractions of *cis* and *trans* azo-PC in the bilayer using the simulation data for the bending rigidity and the membrane thickness, see Fig. 4C. Increasing the *trans* azo-PC fraction in the membrane from 0 to 100 mol % causes a decrease in 1/*β* from 35.6 ± 0.2 to 17.3 ± 0.1. These results indicate stronger interleaflet interactions in the *trans* azo-PC bilayer. The opposite is true for increasing fractions of *cis*-azo-PC, which result in 1/*β* values corresponding to freely sliding monolayers, i.e., interleaflet interactions become weaker. All these results clearly demonstrate that membrane elasticity of azo-PC vesicles and monolayer interactions can be conveniently regulated by light.

### 3.5. Specific membrane capacitance and dielectric constant of lipid vs azo-PC bilayers

The observed changes in the membrane thickness between *cis* and *trans* azo-PC GUVs, prompted us to explore differences in the electrical properties of these membranes, which are thickness dependent such as the membrane capacitance. This property quantifies the ability of a membrane to store electrical charge and determines the propagation velocity of action potentials. The membrane capacitance of a vesicle or cell is proportional to their surface area while the specific membrane capacitance is normalized by the surface area. Thus, the specific membrane allows accurate comparison of vesicles and cells irrespective of their size and shapes.^[76]^ We utilized an approach introduced by Salipante et al.^[38a]^ based on GUV electrodeformation, to deduce the specific membrane capacitance of POPC and azo-PC membranes and subsequently interrogate the membrane dielectric constant (see section 2.8). To the best of our knowledge, this is the first study addressing the specific membrane capacitance and dielectric constant of azo-PC membranes.

Phase contrast micrographs of vesicles were recorded under a frequency sweep of the applied electric field (see Fig. 5A) and used to obtain the GUV aspect ratio (Fig. 5B). The critical frequency at which GUVs undergo prolate-to-oblate transition is used to obtain the specific membrane capacitance (see Eq. 2 and section 2.9).

**Figure 5:**
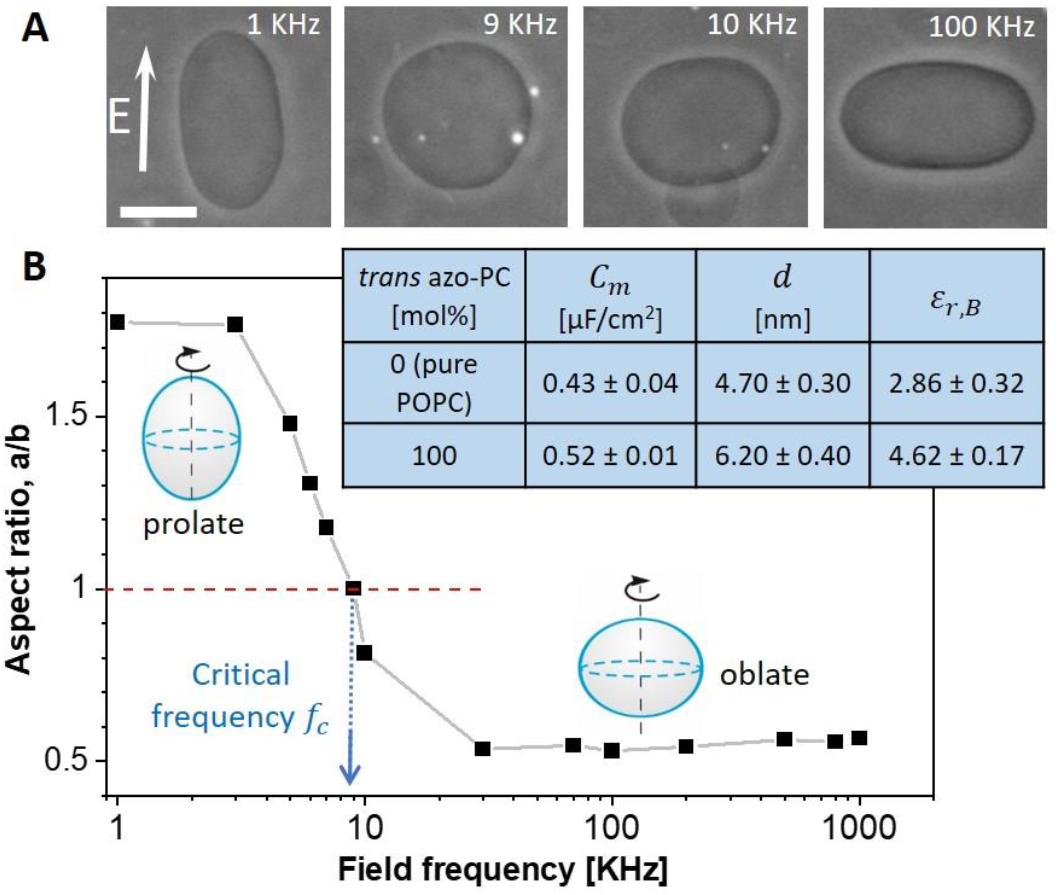
Specific membrane capacitance measurements and estimates of the dielectric permittivity of 100 mol% *trans* azo-PC and pure POPC membranes. (A) Phase-contrast images of a GUV exhibiting morphological prolate-sphere-oblate transitions under a frequency sweep at a field strength of 10 kV/m. The scale bar is 10 μm. (B) Aspect ratio versus frequency plot of the same GUV shown in (A). The critical field frequency (∼9 KHz) of the prolate-oblate transition point with aspect ratio *a/b* = 1 is indicated with an arrow. The table in the inset summarizes the specific membrane capacitance (averaged over 10 vesicles per composition), results for the mean bilayer thickness obtained from AFM (see also Fig. S7), and the estimated dielectric constant for pure POPC membrane and pure azo-PC bilayer in the *trans* state.

The method was applied to measure pure POPC and 100% azo-PC vesicles. We intended to explore the effect of azo-PC isomerization, but because the frequency sweep of a single GUV takes approximately 15-20 minutes, the long-term exposure of the GUVs to UV illumination led to membrane tubulation and complex morphological transitions, which we believe are related to photooxidation of the bilayer components. Furthermore, the deformations prevented us from reliably detecting the transition frequency. Thus, data for membranes in the *cis* state are not included. The table in Fig. 5D summarizes the specific membrane capacitance values of pure POPC and 100 mol% *trans* azo-PC bilayers. The specific membrane capacitance of pure POPC GUVs, 0.43 ± 0.04 μF/cm^2^, is consistent with literature values^[61]^. However, for GUVs composed of pure *trans* azo-PC, the specific membrane capacitance significantly increased to 0.52 ± 0.01 μF/cm^2^.

The specific membrane capacitance is the resultant capacitance of a series of three capacitors: the bare lipid membrane and the adjacent space charge regions in the solution on both sides of the bilayer, see section 2.9. The bare lipid capacitance scales inversely with its thickness (Eq. 4), which is why the larger thickness and capacitance of *trans* azo-PC bilayers compared to pure POPC ones is counterintuitive, unless the membrane dielectric constant changes as well. Using the obtained values for the specific membrane capacitance from vesicle electrodeformation and bilayer thickness from AFM, we deduced the dielectric constants of each membrane. The dielectric constant of the *trans* azo-PC bilayer was found to be 4.62 ± 0.17, almost twice higher than that of POPC, which was found to be 2.86 ± 0.32 (see the table in Fig. 5D).

### 3.6. Photoresponse of minimal cells to exogenous addition of azo-PC

Above we provided a detailed characterization of photo-triggered membrane remodeling events on vesicle bilayers prepared from POPC and varying fractions of azo-PC. In view of the potential application of the photoswitch to modulate the area, morphology, material and electric properties of cellular membranes, we raise the question whether it is possible to observe similar membrane dynamics and photoresponse upon incubating pure POPC membranes in solutions containing the azo-PC photoswitch. Such exogenous incorporation of azo-PC into the already established membrane bilayers is a prerequisite for the direct manipulation of cells. We hypothesized that the amphiphilic nature of azo-PC molecule may enable the membrane to adsorb this photoswitch from the external media to the outer leaflet of the bilayer asymmetrically thus rendering the membrane photoresponsive.

POPC GUVs were prepared and incubated in a solution of azo-PC at a concentration equal to that of the total lipid concentration in the GUV suspension (see Methods section 2.10). In this way, we aimed at obtaining vesicles with 50 mol% azo-PC. After 20 min incubation in the azo-PC solution, and prior to UV illumination, most GUVs appeared to exhibit thick outward protrusions (see first snapshot in Fig. 6A). Controls based on the addition of the same amounts of azo-PC-free solution resulted in no detectable changes in the GUV morphology.

**Figure 6:**
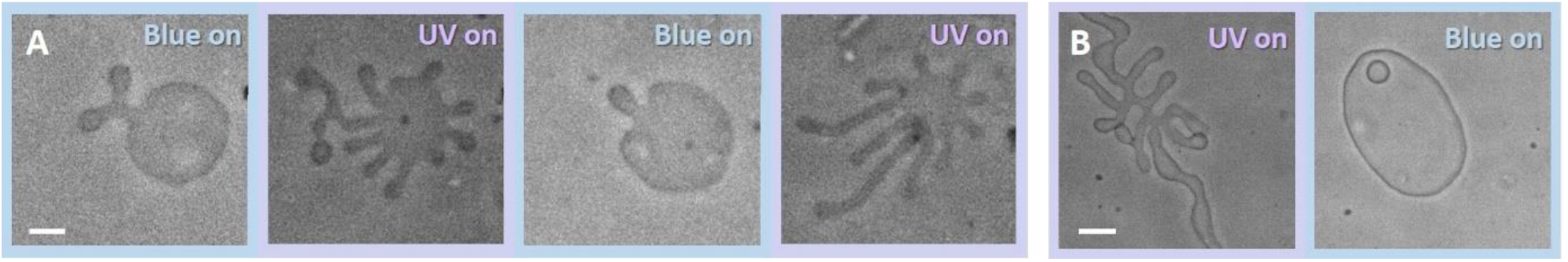
Two examples of the photoresponse of POPC vesicles exogenously doped with azo-PC and irradiated with UV and blue light. The vesicles were exposed to a solution of azo-PC at 15.56 μM bulk concentration (equivalent to the total lipid concentration in the GUV suspension). The GUVs adopt highly tubulated morphologies under the exposure of UV light (365 nm). Under blue light irradiation (450 nm), most of the tubules are re-adsorbed and the vesicles adopt their initial non-tubulated morphology, see also Movie S8. Illumination conditions are indicated on upper-right side of each snapshot. The scale bars correspond to 10 μm.

Upon UV illumination, quasi-spherical GUVs with a few outer protrusions transform into tubular networks or highly tubulated morphologies. This process occurs within a couple of seconds of irradiation (see Movie S8). The protrusions retract back under blue light (Fig. 6). These morphological transformations could be reversed back and forth over multiple photoswitching cycle (Fig. 6A). This result demonstrates the reversibility of the light-induced manipulation of exogenously doped azo-PC vesicles. Thus, we conclude that it is feasible to achieve light-triggered membrane remodeling events on cells exogenously exposed to azo-PC, similarly to exposing GUVs as artificial cells.

### 3.7. Molecular shape and flip-flop free energy of azo-PC

Asymmetric distribution of amphiphilic molecules in the membrane is known to induce membrane tubulation stabilized by spontaneous curvature^[77]^ of the bilayer. The outward formation of relatively thick tubes of micron-sized diameters that we observe (Fig. 6) indicates that the bilayer has only small positive spontaneous curvature, which implies that the final distribution of azo-PC in the GUVs upon exogenous addition is only slightly asymmetric. We speculate that the externally added azo-PC translocate to the inner membrane leaflet either via defects or flip-flop to equilibrate the surface coverage in both leaflets. Azo-PC was introduced in the GUV sample as a solution of dichloromethane/methanol. Before evaporation, the organic solvents could create temporary defects in the membrane allowing material exchange between the leaflets thus balancing the asymmetric distribution of azo-PC.

To explore flip-flop as a possible mechanism, we estimated the energy barrier for translocating azo-PC across the membrane by performing PMF calculations (see section 2.6). Analyzing a bilayer made of 25 mol% azo-PC and 75 mol% POPC allowed us to directly compare the PMF of interleaflet translocation of POPC and of azo-PC when the photoswitch is in *trans* and in *cis* conformation (Fig. 7A,B,D,E). The translocation of POPC in pure POPC bilayer was also examined (see SI Fig. S8). All PMFs for POPC and different conformation of azo-PC showed similar energy barrier of 47-50 kJ/mol for the flip-flop of both molecules across the bilayer. Considering that typical flip-flop times of lipids are on the order of an hour ^[78]^, we find that azo-PC flip-flops at a similar time scale and we can thus exclude flip-flop as a possible mechanism of azo-PC translocating to the inner leaflet upon exogenous addition to GUVs.

**Figure 7:**
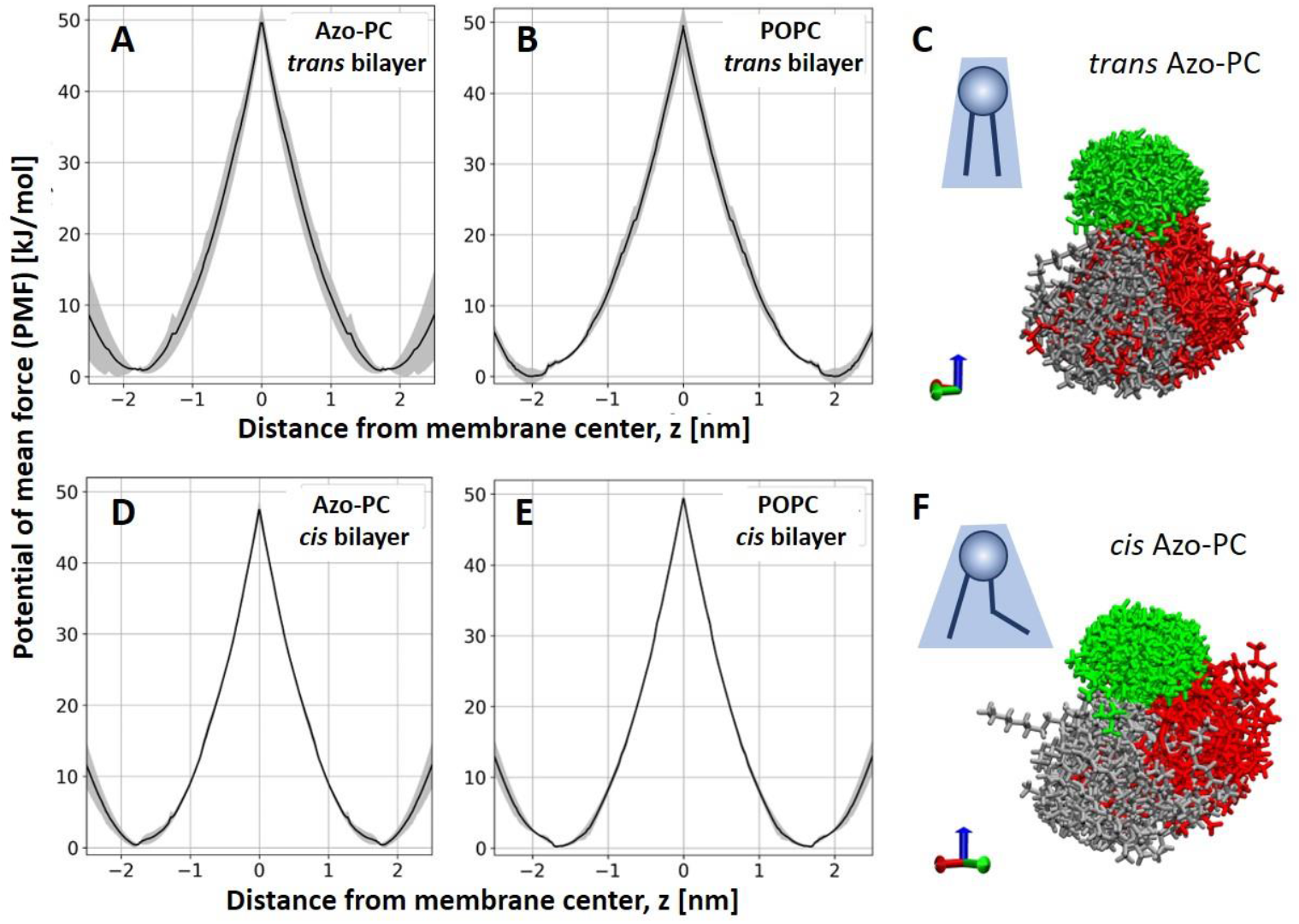
Free energy of flip-flop of azo-PC and molecular curvature. (A, B, D, E) PMF calculation plots for the flip-flop energy of azo-PC (A, D) and POPC (B, E) in a bilayer containing 25 mol% azo-PC and 75 mol% POPC when the photoswitch is in the *trans* (A, B) or in the *cis* state (D, E). The maxima of the plots illustrate the energy barrier for flip flop. Standard error of each calculation is shown in grey. (C, F) Snapshots 100 aligned and centered azo-PC molecules (see text for details) in *trans* (C) and *cis* (F) state. Green, gray and red corresponded to PC head groups, hydrocarbon tails and azobenzene tails, respectively. The blue arrow on the bottom right indicate the direction of the membrane normal.

We also explored the molecular curvature of azo-PC, comparing the *trans* and *cis* states. For this, we generated MD simulation snapshots from 100 azo-PC molecules aligned at the C2 carbon (the atom where the two tails come together) rotating the molecules around the *z* axis to align the first two bonds along each chain (see Fig. 7C,F). This presentation produces an apparent spatial cloud explored by an azo-PC molecule in the bilayer, giving a rough idea of the molecular curvature of azo-PC in POPC environment. The results illustrate that in the *cis* state the azo-PC hydrophobic tails explore or occupy a larger volume (similarly to phosphatidylethanolamine lipids) compared to the molecular arrangements of the molecule in the *trans* state, suggesting that *cis* azo-PC may generate higher curvature compared to *trans* azo-PC.

## 4. Discussion

Our results demonstrate that light can be employed as a facile, fast and sustainable tool for micromanipulation of artificial minimal cells in combination with the photoswitch azo-PC as an efficient converter of light to mechanical energy. As mentioned in the introduction, several other studies have investigated the response of azo-PC membranes to light. As we will discuss below, we find that some of the previously published data are inconsistent among themselves (even though reported from the same group) and partly with our and will propose reasons to explain these inconsistencies.

### 4.1. Membrane area changes upon isomerization

Our results reveal that photoisomerization of azo-PC under UV and blue light triggers complex shape transformations and budding events in the membrane (Fig. 1C,D) also reported in refs. ^[32c, 34a-c]^ for various photoswitches. Here, using several model membranes, we show that area increase as much as 20 % can be obtained for pure azo-PC membranes (Fig. 2E). Our results clearly demonstrate the deficiencies of measuring area changes using DLS on LUVs for this purpose. Indeed, previous measurements suggested very small changes in the diameter of pure azo-PC LUVs of the order of 3 %^[34a]^ implying area change of only ∼6 % which is obviously unrealistic considering the directly observed area change seen with microscopy of GUVs and accompanying simulations. Similarly, photoinduced size changes of SUVs made of an azo-PC derivative showed tiny increase of 1 %^[63, 79]^, which could be due to the fact that the measurement is model dependent (X-ray scattering) but also because the SUV membrane is highly curved.

We also showed that the membrane area can be finely tuned by altering the molar fraction of azo-PC in the membrane thus modulating shape-dependent cellular processes such as endo/exocytosis and intra/intercellular trafficking. The potential phototoxicity of UV and blue light can be remedied by shifting the excitation wavelength of azobenzene derivatives into the lower energy range as recently demonstrated^[79]^. Coupling optical microscopy and GUV electrodeformation allowed us to precisely quantify the reversibility and kinetics of the photoisomerization process over the whole membrane. These measurements are also direct (not model dependent) and just like seeing is believing, we consider them reliable. Among previously published data, this is also the first report to directly and accurately quantify the dose dependent, reversible area changes of membranes containing azo-PC photolipids.

### 4.2. Membrane bending rigidity and leaflet coupling

We find inconsistencies in the literature also regarding the membrane mechanical properties, in particular, the bending rigidity. Using optical tweezers and a model for the deformation of GUV trapped locally and deformed by a flow, Pernpentier et al.^[34a]^ reported bending rigidity values for *cis* and *trans* azo-PC GUVs, κ_*cis*_ = (5.4 ± 1.8) × 10^−19^ J and κ_*trans*_ = (1 ± 0.6) × 10^−17^ J corresponding to 131 k_B_T and 2433 k_B_T, respectively. First, such bending rigidities, and especially that of *trans* azo-PC would correspond to very stiff membranes (like gel phase)^[80]^, which is inconsistent with the fact that these membranes are liquid disorder in nature. Second, at such high bending rigidity the membrane should not fluctuate, which is also not the case in their study, suggesting that the model used to calculate the values ^[34a]^ is incorrect (presumably, the assumption for ellipsoidal deformation affects the calculation).

Another study by the same research group^[34c]^, reports the following values for *cis* and *trans* azo-PC membranes measured micropipette aspiration of GUVs, κ_*cis*_ = 6.4 × 10^−21^ J and κ_*trans*_ = 3.1 × 10^−20^ J (respectively corresponding to 1.6 k_B_T and 7.5 k_B_T), largely contradicting their own previous results. The reason for these extremely low values, could be that micropipette aspiration measurements rely on the mechanical deformation of the membrane and are known to suffer from stretching elasticity contributions^[70]^ as demonstrated in more detail by Henriksen and Ipsen^[81]^. In this respect, fluctuation spectroscopy, not relying on mechanical perturbation of the membrane is the gold standard, providing us with more reliable results; this is supported also by the agreement with literature data for the bending rigidity of pure POPC bilayers^[70]^.

Our experimental results for azo-PC dose-dependent bending rigidity and area increase are fully consistent with MD simulations (Figs. 2E and 4A), especially for azo-PC fractions smaller or equal to 50%. Using this as evidence, we explored additional elastic parameters, not accessed experimentally, namely the intermonolayer coupling which hasn’t been characterized previously.

The coupling constant 1/β of *trans* azo-PC membrane is close to values reported for 1,2-dioleoyl-sn-glycero-3-phosphocholine (DOPC), 1/β_DOPC_ ≈ 18^[72a]^. For POPC we find 1/β ≈ 36. DOPC with its two unsaturated long chains of (18:1) oleic acid, must form a relatively tightly coupled bilayer compared to POPC with only one unsaturated chain. *Trans* azo-PC significantly increases the interleaflet interactions monolayers compared to *cis* azo-PC, which decreases the coupling, showing a similar trend as that reported for a fusion peptide^[72a]^ that was shown to shift the coupling towards freely sliding monolayers. The weaker interleaflet interactions for *cis* azo-PC containing membranes may be caused by reorientation of the azobenzene tail somewhat parallel to the membrane plane reducing the coupling and facilitating interleaflet slip. This idea is corroborated by the spatial distribution and molecular occupancy that we find from MD simulations for the *cis* state as shown in Fig. 7F.

### 4.3. Electrical properties and membrane thickness

We characterized the electrical properties of the azo-PC membranes providing the first values for the specific membrane capacitance and the dielectric constant of azo-PC bilayers. In our studies, the specific membrane capacitance, and the thickness of azo-PC and POPC bilayers obtained from separate experiments (GUV electrodeformation and AFM on membrane patches, respectively) presented higher values for membranes with *trans* azo-PC compared to pure POPC bilayers. Previously, results on increasing membrane capacitance have been reported for an amphiphilic photoswitch (Ziapin2) in neuronal membranes^[82]^, but have been attributed to membrane thinning.. We refrain from comparing these data to ours on specific membrane capacitance (which is the area-rescales capacitance) as the area changes related to insertion of Ziapin2 in the membrane are not known. We interpret our observations for the larger specific capacitance and thickness of the *trans* azo-PC membrane compared to POPC (table in Fig. 5) as resulting from higher dielectric constant signifying higher ability to store electrical energy and to polarize in electric fields.

Our AFM measurements showed a dramatic thickness decrease of ∼1.5 ± 0.6 nm of the azo-PC bilayer upon *trans*-to-*cis* isomerization (Fig. SB). This is consistent with the decrease in the hydrophobic thickness as observed with MD simulations (Fig. 4B). Membrane thickness data for azo-PC membranes from X-ray scattering on azo-PC SUV suspensions reported only a 0.4 nm decrease in the membrane thickness associated with the *trans-cis* isomerization. Considering that the membrane as an elastic sheet should preserve its volume (equal to the area times the thickness) upon thinning, we believe that our results are much more realistic as an area increase by 20-30 % (Fig. 2E) would roughly correspond to 17-24% decrease in the membrane thickness, which is of the order of 1.1-1.5 nm as seen by the AFM measurements.

From a structural point of view, we note that azo-PC in the trans conformation partly resembles the lipid DSPC (1,2-distearoyl-sn-glycero-3-phosphocholine), having the same fatty acid tail. The DSPC bilayer thickness is approximately 5.8 nm and 5.5 ± 0.5 nm^[83]^.

In conclusion, through careful integration of accurate experimental and computational methodologies, we addressed an existing knowledge gap by delivering a comprehensive depiction of the membrane response to the photoisomerization of azo-PC across a range of model membrane systems. We critically evaluated current reports and existing discrepancies providing an encompassing understanding of membrane expansion, thinning, softening, interleaflet coupling, capacitance and dielectric constant. We should also mention that we cannot exclude that some of the above-mentioned issues with discrepancies in the literature could be associated with the specific source of azo-PC lipids used in the different studies (commercial as here vs synthesized by other authors) and, potentially, the different fraction of impurities.

Overall, our studies with GUVs, LUVs, SLBs, Langmuir monolayers and MD simulations consistently illustrated that light-induced membrane deformations due to photoisomerization of azo-PC cause dynamic alterations in the membrane. These include changes in bilayer packing, membrane elasticity as well as interleaflet interactions thus leading to dramatic changes in the material and electric properties of membranes, which scale with azo-PC fraction. Lastly, the reproducible photoresponse of already formed vesicles after the exogenous insertion of azo-PC in the membrane provides a promising background for intercalating this photoswitch in more complex cell systems to optically control cellular activities.

## Author contributions

MA performed all vesicle-based work. AG performed the simulations. FC performed the experiments on monolayers. MA, VG, FC, AG analyzed data. NY prepared SLBs and generated AFM data. SB analyzed AFM data. RD and JH proposed the project. RD supervised the project. MA and RD wrote the manuscript with contributions from all authors.

## Acknowledgements

MA acknowledges funding from the International Max Planck Research School on Multiscale BioSystems. RD and MA thank M. Miettinen for the fruitful discussions on the stability and flip-flop in asymmetric membranes. MA thanks V. Vitkova for the feedback on the calculations of the dielectric constant of vesicles. This work was supported by Germany’s Excellence Strategy, EXC 2008/1 (UniSysCat), Grant 390540038.

