## Supporting information for "Photomanipulation of minimal synthetic cells: area increase, softening and interleaflet coupling of membrane models doped with azobenzene-lipid photoswitches"

### Modulating membrane shape and mechanics of protocells by light: area increase and softening of membrane models doped with azobenzene-lipid photoswitches

Mina Aleksanyan[1,2], Andrea Grafmüller[1], Fucsia Crea[3], Vasil N. Georgiev[1], Naresh Yandrapalli[1], Stephan Block[2], Joachim Heberle[3], Rumiana Dimova[1]*

[1] Max Planck Institute of Colloids and Interfaces, Science Park Golm, 14476 Potsdam, Germany

[2] Institute for Chemistry and Biochemistry, Freie Universität Berlin, Berlin, Germany;

[3] Department of Physics, Freie Universität Berlin, Berlin, Germany

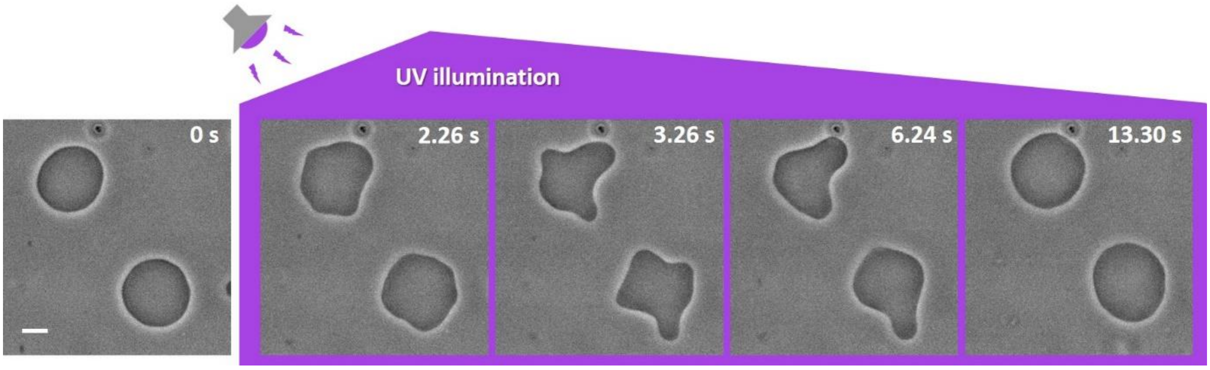


**Figure S1:** Time sequence of the response of GUVs made of azo-PC:POPC 50:50 mol% to *trans*-to-*cis* photoisomerization under UV illumination, see also Movie S3. The GUVs were grown in 100 mM sucrose solution and 1:1 diluted in 105 mM glucose solution. The time stamps show the time after starting the UV irradiation. The scale bar corresponds to 10 µm. UV illumination triggers complex shape transformations in GUVs over time.

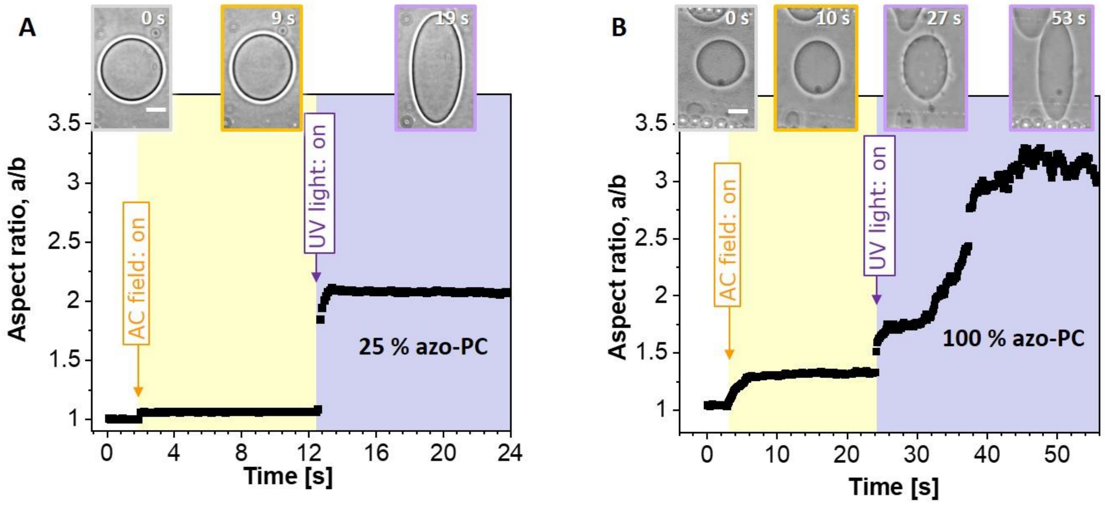


**Figure S2:** Deformation of vesicles composed of 25 mol% (A) and 100 mol% azo-PC (B) under the influence of UV light. The graph shows the vesicle degree of deformation (a/b, aspect ratio) over time. The snapshots illustrate vesicle shape changes over time. The color code is as follows: Gray frames correspond to the vesicles in the absence of AC-field and UV-light, orange frames correspond to the vesicles exposed to AC-field (5kV $m^{-1}$ and 1 MHz), purple frames correspond to vesicles exposed to both AC-field (5kV $m^{-1}$ and 1 MHz) and UV illumination (365 nm). The scale bar corresponds to 20 µm. As azo-PC fraction in the membrane increases from 25 to 100 mol %, GUVs undergo larger membrane deformations through complex shape transformations and budding events slowing down vesicle response and increasing the time for the GUV to reach to the maximum degree of deformation.

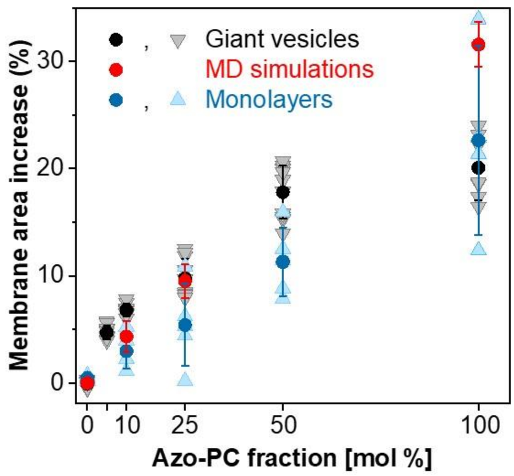


**Figure S3:** Individual measurement data points for membrane area expansion associated with photoisomerization as assessed from GUV electrodeformation (gray symbols show data on individual GUVs; black data given as mean values +/-SD are as in Fig. 2E in the main text) and monolayers (light blue triangles show measurement of individual Langmuir monolayers, dark blue – mean +/- SD as in Fig. 3E). The MD simulations data (red, as in Fig. 3E) are also included.

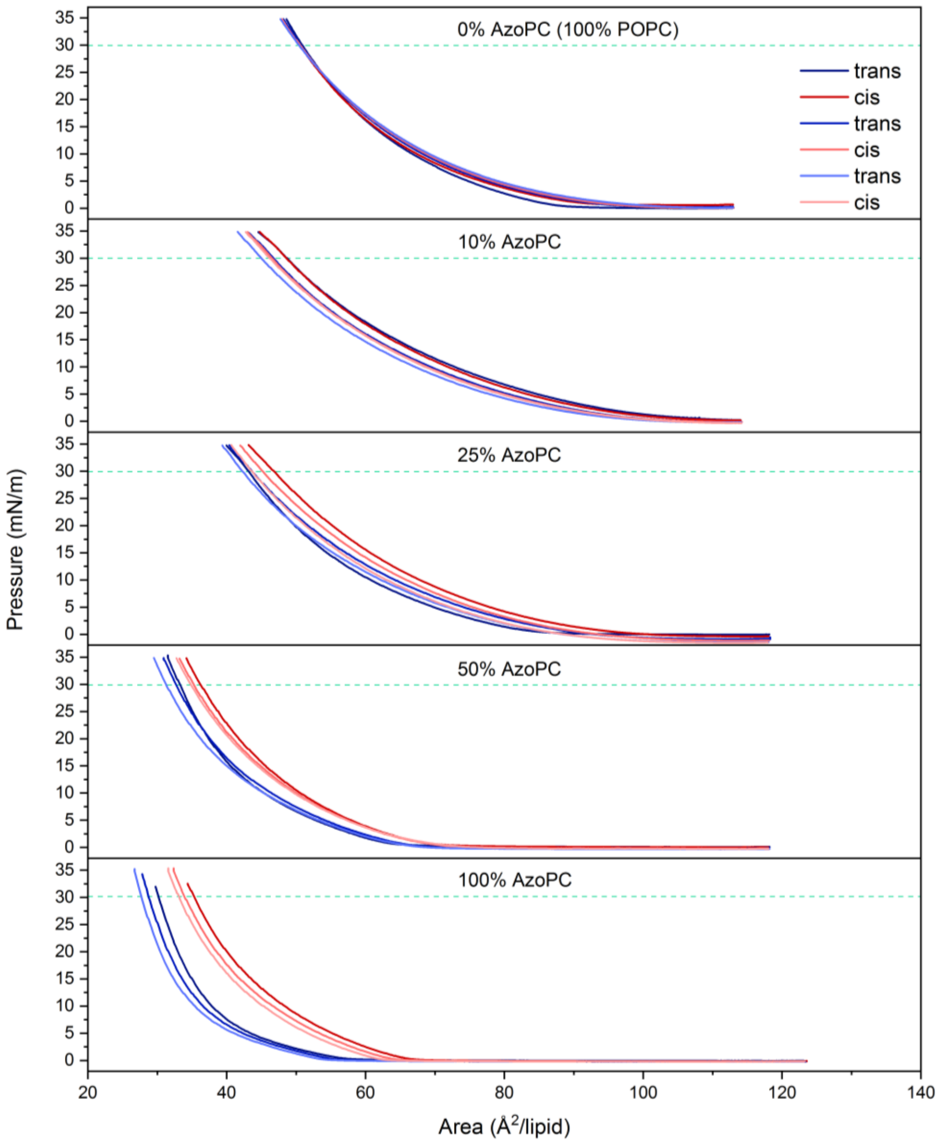


**Figure S4:** Example pressure-area isotherms of lipid monolayers containing different fractions of azo-PC measured consecutively. The area values at 30 mN/m pressure (indicated with a green dashed line in all panels) were used for calculating the monolayer area expansion reported in Fig. 2E in the main text. The small drift towards smaller area between consecutive isotherms of monolayers with the same composition are due to lipid loss occurring in cycles of compression isotherms. The pairs of isotherms considered for the calculation of the area expansion are always two consecutive ones, one *trans* and one *cis*. For visual clarity, only one set of isotherms per film composition is shown. The area increase of the films containing 10 % (mol) Azo-PC was on average 1.3 ± 1.1 % (standard deviation); for the 25 % film it was 5.0 ± 2.2 %; for the 50 % film it was 10.9 ± 3.8 %; for the 100 % it was 19.5 ± 6.4 %. The control experiment with a pure POPC monolayer showed no expansion (-0.20 ± 0.38 %). With increasing Azo-PC fraction in the lipid monolayers the pressure-area isotherms show larger expansion.

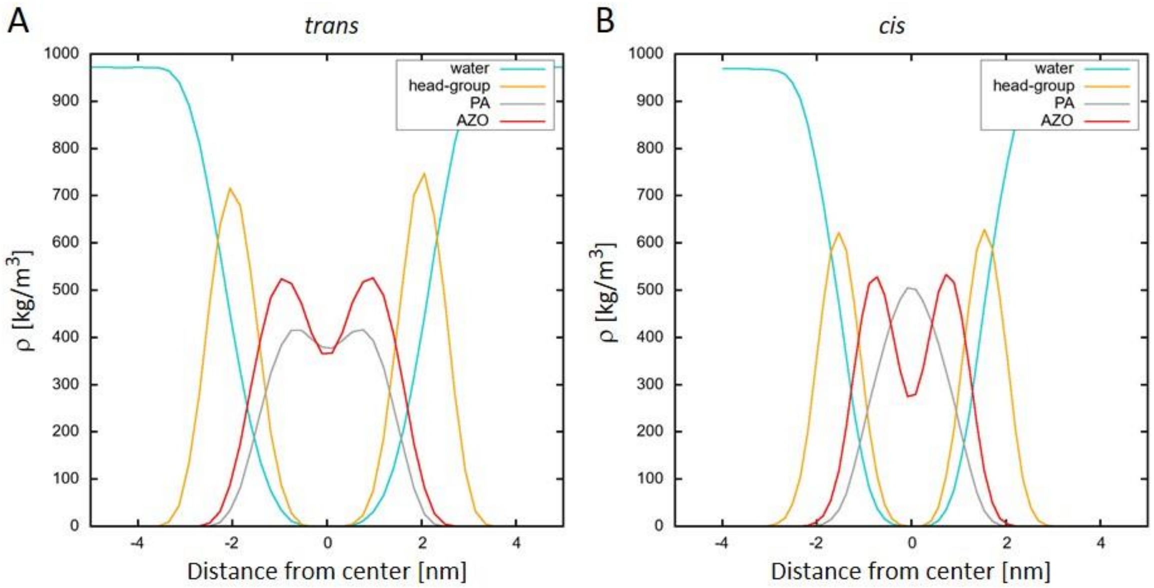


**Figure S5:** Radial density profiles from MD simulations of 100 mol% azo-PC in (A) *trans* and (B) *cis* conformation. The conformational change from *trans* to *cis* of azo-PC (AZO, red) leads to relocation of the palmitoyl chains (PA, gray) deeper in the bilayer and the headgroup region becomes better hydrated.

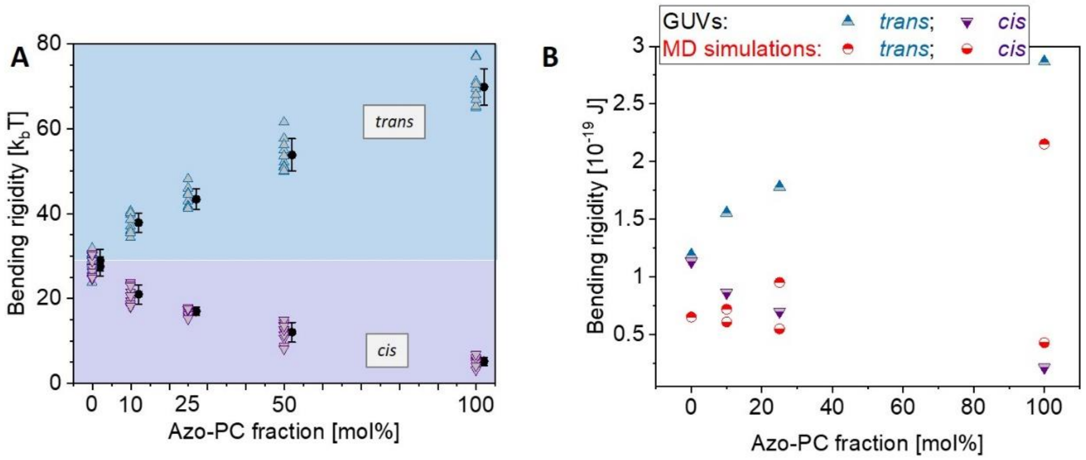


**Figure S6:** Raw data for the values of bending rigidity of POPC membranes with various azo-PC fractions obtained from experiments and simulations. (A) Experimental data obtained from fluctuation spectroscopy: open triangles show measurements on individual vesicles; solid circles and error bars indicate mean and standard deviations. (B) Mean values of the bending rigidity (in Joules) obtained from experiment (triangles – same mean data +/-SD as in B) and MD simulations (circles). As *trans* azo-PC fraction in the membrane increases, GUVs become stiffer. As *cis* azo-PC fraction in the membrane increases, GUVs become softer.

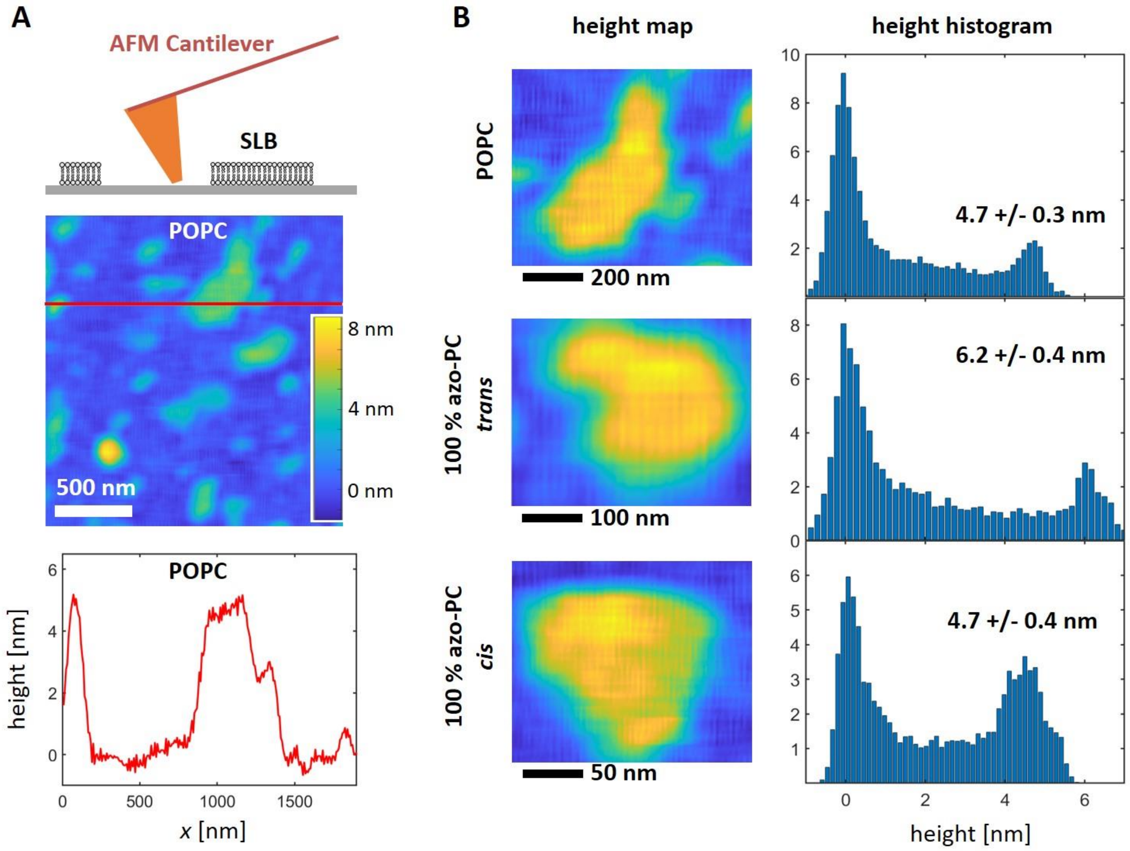


**Figure S7:** Thickness measurements of POPC and 100 % azo-PC supported lipid bilayers (SLBs) using atomic force microscopy (AFM). (A, top) Schematic representation of the measurement geometry, in which SLB patches were formed on glass surfaces and imaged using liquid phase AFM. (A, middle) Representative height map of a sample containing POPC patches and (A, bottom) a typical cross section in the height map. (B, left) AFM screenshots of example patches of pure POPC (top) and azo-PC SLBs in *trans* (middle) and *cis* state (bottom), and (B, right) the corresponding height histograms. The thickness of the bilayers made of POPC, 100 mol % *cis* and *trans* azo-PC measured as 4.7 ± 0.3, 4.7 ± 0.4 and 6.2 ± 0.4 nm, respectively.

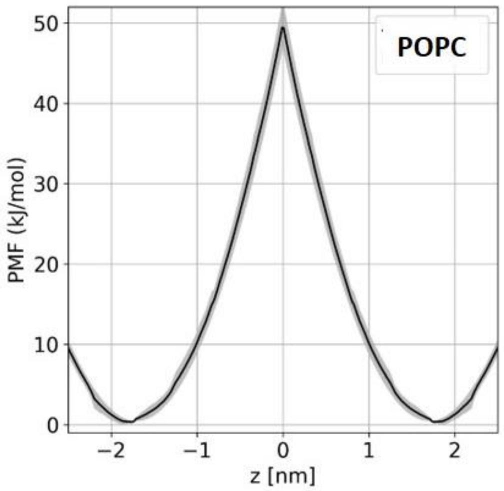


**Figure S8:** Potential mean force calculations (PMF) of interleaflet translocation of a POPC lipid in POPC bilayer. The potential maximum of the plot illustrates the energy barrier for flip flop while ‘z’ is the displacement coordinate being perpendicular to the lipid bilayer. Standard error of the calculation is shown in grey. PMF profiles of POPC, *cis* and *trans* azo-PC lipids showed the same trend indicating the absence of flip-flop in the experimental time scales for all these lipid compositions.

**Movie captions**

**Movie S1:** A sequence of confocal images of a sample of GUVs made of 100 mol% azo-PC labelled with 0.1 mol% Atto-647N-DOPE monitored during photoisomerization (images are acquired roughly every 1.3 s). Time stamps show time after initiating acquisition. The initial radius of the larger vesicle is approximately 20 µm. The frames in which UV irradiation is applied are marked in the upper part of the images. The video was processed with LAS X, Fiji and HandBrake software.

**Movie S2:** Real time video of 50 mol% azo-PC GUV acquired under phase contrast at 50 frames per second (fps); same vesicle as in Fig. 1D in the main text. The initial radius of the GUV is approximately 12 µm. The time stamps and the period of UV irradiation are indicated in the upper part of the images. The video was processed with FiJi and HandBrake software.

**Movie S3:** Real time video (50 fps) of two 50 mol% azo-PC GUVs shown also in Fig. S1. The radii of GUVs are approximately 10 and 12 µm. The time stamps and the period of UV irradiation are indicated in the upper part of the images. The video was processed with FiJi and HandBrake software.

**Movie S4:** Phase contrast video (8 fps) of a POPC GUV during electrodeformation and UV irradiation; the data from this vesicle is plotted in Fig. 2B. The AC-field amplitude is 5 kV $m^{-1}$ at a frequency of 1 MHz. The initial GUV radius is approximately 14 µm. The time stamps, electric field application and UV illumination are indicated in the upper part of the images. The video was processed with FiJi and HandBrake software.

**Movie S5:** Phase contrast video (8 fps) of a 10 mol% azo-PC GUV during electrodeformation (5kV $m^{-1}$, 1 MHz) and UV irradiation; the data from this vesicle is plotted in Fig. 2B. The initial GUV radius is approximately 25 µm. The time stamps, electric field application and UV illumination are indicated in the upper part of the images. The video was processed with FiJi and HandBrake software.

**Movie S6:** Phase contrast video (8 fps) of a 50 mol% azo-PC GUV during electrodeformation (5kV $m^{-1}$, 1 MHz) and UV irradiation; the data from this vesicle is plotted in Fig. 2C. The initial GUV radius is approximately 15 µm. The time stamps, electric field application and UV illumination are indicated in the upper part of the images. The video was processed with FiJi and HandBrake software.

**Movie S7:** Phase contrast video (16 fps) of a 10 mol% azo-PC GUV under AC field (5kV $m^{-1}$ and 1 MHz) and UV and blue-light illumination, illustrating reversible photoswitching over multiple cycles; data from this vesicle is plotted in Fig. 3B. The initial GUV radius is approximately 11 µm. The time stamps, electric field application, UV and blue light illumination are indicated in the upper part of the images. The video was processed with FiJi and HandBrake software.

**Movie S8:** Phase contrast video (acquired at 100 fps) of POPC vesicles exogenously doped with azo-PC and irradiated with UV and blue light; the larger vesicle is shown in Fig. 7A. The time stamps, UV and blue light illumination are indicated in the upper part of the images. The video was processed with FiJi and HandBrake software.
